# Mining association rules for targeted spatiotemporal aquatic environmental DNA (eDNA) sampling

**DOI:** 10.64898/2026.09.16.752056

**Authors:** Nikolett Toth, Luiza Antonie, Robert H. Hanner, Daniel J. Gillis, Jarrett D. Phillips

**Author notes:** **Corresponding Author**: Jarrett D. Phillips^1,3^, **Email Address**.

## Abstract

Environmental DNA (eDNA) offers a non-invasive alternative to traditional, more destructive sampling methods for determining species occupancy at ecological sites of interest. Aquatic eDNA sampling entails filtering a known volume of water to capture and detect genetic material shed by organisms. While the influence of individual environmental properties on the presence of target eDNA has been widely studied, it remains unclear how variables like temperature, pH, flow rate and conductivity correlate collectively with site electrofishing counts and eDNA concentrations. Resolving this question is important for two reasons: (1) typically, only eDNA, not physical specimens, is collected and measured, and (2) when methods like electrofishing and eDNA sampling are used in tandem, results often differ. Here unsupervised association rule-based machine learning is employed to discover interesting relationships among sampled covariates within a previously published case study of native brook trout (*Salvelinus fontinalis*) collected from Hanlon Creek (Guelph, Ontario, Canada) in September 2019. From a dataset of only 126 observations, the mining process revealed over 12000 plausible association rules linking covariates to eDNA concentrations (low/high) and electrofishing outcomes (absence/presence of brook trout). A strict pruning strategy reduced this ruleset to a manageable size of 153 associations, some of which were corroborated by existing literature, and some of which were novel (such as those potentially relating electrical conductivity to microbial and enzymatic activity). The entire workflow is included as a new R package called `RulesTools`. These results highlight the promise of association rule mining as a tool for guiding eDNA metadata collection, complementing statistical modelling, and informing conservation and management decision-making.

**Open Research Statement:** The `RulesTools` R package is available for download through the Comprehensive R Archive Network (CRAN) at https://cran.r-project.org/web/packages/RulesTools/index.html and via GitHub at https://github.com/nikolett0203/RulesTools. The brook trout dataset utilized in this study can also be found at the above links.

## 1 Introduction

Biodiversity is in crisis as many native species face the threat of extinction within the coming decades and invasive species rapidly expand their ranges (Keck et al., 2025). As such, novel means of assessing species-level ecosystem impacts are increasingly necessary. Environmental DNA (eDNA) (Ficetola et al., 2008) has emerged as a promising technique to quantify the extent of taxon presence or absence at ecological sites of interest. It involves the collection of genetic material (such as epithelial cells or metabolized excreta) via water, soil, sediment, or air sampling. Unlike traditional capture-based sampling techniques (*e.g.* gillnetting, electrofishing), eDNA sampling provides a non-invasive, highly sensitive, and often more convenient means of monitoring ecosystem health. However, widespread adoption of this technique requires an understanding of myriad abiotic factors that regulate the origin, state, transport, and fate of eDNA and thereby affect species distribution patterns (Barnes and Turner, 2016).

eDNA persistence in aquatic environments is driven by diverse physicochemical influences including temperature, pH, dissolved oxygen levels, flow rate, and electrical conductivity. Several recent studies have examined these associations and their impact on species detection and occupancy. Through detailed microcosm experiments, Strickler et al. (2015) assessed the role of UV-B radiation, temperature, and pH on American bullfrog (*Lithobates catesbeianus*) tadpole eDNA degradation. Findings indicate that temperature is negatively correlated, and pH positively correlated, with detectable eDNA concentrations. That is, high temperatures hinder eDNA persistence (McCartin et al., 2022) while alkaline environments maintain it (Seymour et al., 2018). Other studies have noted similar trends among abiotic and biotic conditions within a number of aquatic species: Northern pike (*Esox lucius*) (Ogonowski et al., 2023); Chinese white shrimp (*Fenneropenaeus chinensis*) (Qian et al., 2022); and Blanding’s turtle (*Emydoidea blandingii*), chain pickerel (*Esox niger*), and smallmouth bass (*Micropterus dolomieu*) (Loeza-Quintana et al., 2020). A major criticism of past studies is that they have been conducted *ex situ* as opposed to in natural settings. Furthermore, adequate reporting of abiotic metadata alongside collected environmental samples is severely lacking (Nicholson et al., 2020). Previous work has been conducted to elucidate the impact of environmental variables on the fate of eDNA, leading to the development of five best practices and modelling recommendations for broad adoption by researchers (Harrison et al. 2019). In particular, Recommendation 4 is an express call for the inclusion of environmental metadata alongside water samples to facilitate comparative assessments of eDNA dynamics. Despite this, little work has been accomplished linking the combined effect of physical and chemical attributes on eDNA properties.

eDNA is typically collected by means of portable eDNA samplers (“backpacks”). Two widely available commercial units are the Halltech OSMOS (Halltech Environmental and Aquatic Research Inc.) and the Smith Root (Smith Root Inc.), herein referred to fittingly as “ANDe” (eDNA spelled backwards). Due to eDNA’s contamination risks and its inability to provide direct estimates of species abundance, sampling results are sometimes corroborated by traditional methods such as electrofishing or gillnetting. Despite this, species occupancy gleaned from electrofishing and targeted detection revealed by eDNA sampling are often discordant.

Machine learning approaches (*e.g.* artificial neural networks, random forests, and support vector machines) have recently propelled eDNA research forward, addressing key questions in targeted species detection and metabarcoding across both freshwater and marine ecosystems. However, the majority of published studies and software tools have relied on supervised classification, which requires access to large labelled datasets for training, validation, and testing in order to generalize well to unseen data (*e.g*., Cordier et al. (2018); Dully et al. (2021); Fan et al. (2020); Flück et al. (2022); Keck et al. (2023); Kronenberger et al. (2022)). This is problematic for eDNA research because sample acquisition protocols are not yet standardized (Takahashi et al., 2025). Many important environmental metadata variables are not adequately recorded or are missing altogether within field collection data sheets. Moreover, no studies have yet incorporated unsupervised learning methods within eDNA settings.

Association rules are a form of unsupervised machine learning used for data mining to discover hidden relationships among covariates (Agrawal et al., 1993). Here, this technique is applied as a proof of concept in the context of eDNA sampling, linking abiotic and biotic variables to the detection of native brook trout (*Salvelinus fontinalis*) in Hanlon Creek, Guelph, ON., Canada, based on a previously published study involving electrofishing and eDNA collection in September 2019 (Nolan et al., 2023).

## 2 Methods

### 2.1 Association Rules

Association rules were first introduced in the context of market basket analysis—analyzing supermarket purchases to efficiently discover general but interesting patterns among a set of discretized (categorical, usually binarized) items found within large, sparse, high-dimensional databases (Agrawal et al., 1993). In supermarkets, information gleaned from such rules could be employed to optimally stock shelves or help inform marketing strategies to increase the sale of specific items based on buyer behaviour. Since then, association rule-based mining has seen myriad real-world applications, from analyzing topics in large text corpora (Antonie and Zaïane, 2002), to examining university course enrolments among students (McNicholas, 2006), and identifying differentially expressed genes in cancer microarrays (Antonie and Bessonov, 2011, 2012). More broadly, while association rule mining has seen wide application within bioinformatics and computational biology (Naulaerts et al., 2015; Ceddia et al., 2020; Agostinetto et al., 2022), it has seen little use in ecology (Aloisi et al., 2025). Association rules bring many benefits over more sophisticated statistical models, particularly their ease of understandability and interpretability, as they are inherently nonparametric and data-driven. The association rule mining problem is described as follows. Let *I* be a non-empty set of items, and let *D* be a database consisting of transactions, *T*, where each transaction t, having a unique ID, is a subset of *I* (*i.e.*, *t* ⊆ *I*). A transaction reflects a row in *D* and represents a collection of items that appear together in a single instance, such as products purchased together in a shopping cart or attributes co-occurring in a dataset. Association rules comprise logical implications of the form *X* ⇒ *Y* (read as “*X* implies *Y* “ or equivalently, “if *X*, then *Y* “), where *X* ⊆ *I*, *Y* ⊆ *I*, and *X* ∩ *Y* = ∅. That is, *X* and *Y* are both non-empty subsets of *I* with no common items. In this context, *X* is called the antecedent or left hand side (LHS) and *Y* is termed the consequent or right hand side (RHS). The antecedent usually comprises a logical conjunction (*a* ∧ *b* ∧ …) of items found within it, whereas the consequent typically contains only a single item. *Positive association rules* consider the presence of items in transactions, while *negative association rules* consider items absent from transactions (Antonie and Zaïane, 2004; Antonie et al., 2014). In this study, only positive association rules are considered due the sheer search space added by negative rules.

The overarching goal of association rule mining is to find interesting dependencies among items in datasets of interest. However, the association rule mining task often results in the generation of far too many rules to be practically considered. Given *k* unique items in a transactional database, there are

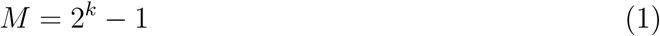

non-empty itemsets in total. Similarly, there are

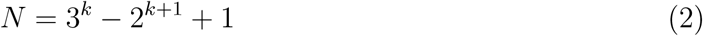

total rules. For example, when *k* = 10, there are *M* = 1023 generated itemsets and *N* = 57002 resulting rules. Often, such rules are trivial, redundant, or both, and hence not useful. Thus, it becomes necessary to prune mined rules, including only the most interesting and actionable associations. Several quantitative measures have been proposed to accomplish this goal. Strong association rules are those that consist of frequently occurring itemsets, in addition to satisfying minimum thresholds for rule support and confidence.

The support of an association rule, Supp(*X* ⇒ *Y*), is the proportion of transactions containing all items from both *X* and *Y* . Mathematically,

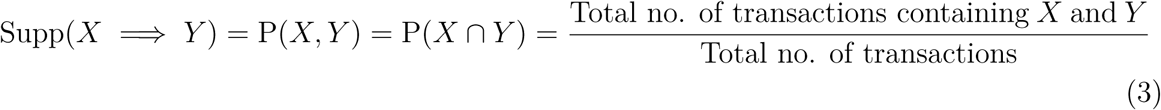

and can be thought of as an estimate of the joint probability of itemsets *X* and *Y* . Note that rules formed from the same items will have identical support. The confidence of a rule, denoted Conf(*X* ⇒ *Y*), is the proportion of transactions involving *X* which also include *Y* . Confidence is analogous to an estimate of the conditional probability of *Y* given *X*. That is,

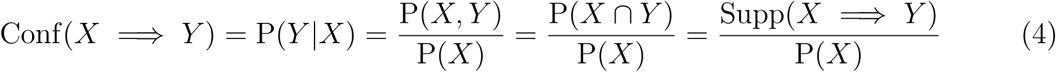

where P(*X*) is the support of itemset *X*. Taken together, support and confidence form the *support-confidence framework* for the identification of interesting and strong association rules.

Establishing appropriate cutoffs for minimum support and minimum confidence is important for several reasons. First, setting these limits too low increases the computational burden and generates an overwhelming number of rules. Many of these rules will be spurious, weak, or noisy, making them difficult to interpret as they often include rare itemsets that lack statistical significance or practical relevance. Low thresholds can also lead to overfitting, where generated rules are too specific and do not extend well to other datasets. In contrast, while higher minimums produce fewer rules, they can also lead to underfitting, whereby rare but meaningful itemsets are filtered out.

Relying solely on support and confidence to rank rules by interestingness is often insufficient, particularly because the confidence of a rule does not consider the support of the consequent, Supp(*Y*). To address this limitation, lift is widely used as an accompanying interestingness measure (Brin et al., 1997). The lift of a rule is given by

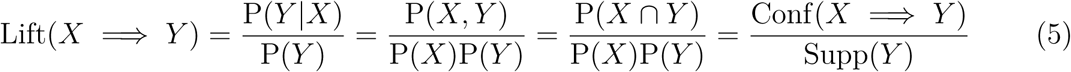

and quantifies the extent to which *X* and *Y* are dependant, where P(*Y*) = Supp(*Y*) is the expected confidence of the rule. A lift value less than one indicates the antecedent and consequent are negatively correlated, whereas a value above one suggests *X* and *Y* are positively related. A value of exactly one signifies no association (*i.e*, independence). In addition to the above-mentioned desirable properties, lift is also symmetric

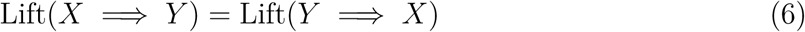

which greatly facilitates the ranking of rules when either or both *X* ⇒ *Y* and *Y* ⇒ *X* or P(*Y* |*X*) and P(*X*|*Y*) may be of interest. Despite this, reliance on raw interestingness measures for ranking and selection of rules is problematic because such measures lack standardization, meaning their ranges are not sufficiently bounded. Numerous attempts have been made to resolve this issue (McNicholas et al., 2008; Shaikh et al., 2018). In the current study, only unnormalized metrics are considered.

### 2.2 Case Study: Brook Trout (*Salvelinus fontinalis*)

#### 2.2.1 Brief Overview

Nolan et al. (2023) compared the efficacy of electrofishing and eDNA capture for detecting native brook trout in Hanlon Creek, Guelph, ON., Canada. The study was conducted in partnership with the non-governmental organization (NGO) Trout Unlimited Canada (TUC), aimed at the conservation, protection, and restoration of freshwaters across Canada. Industry collaborators also participated, including Natural Resources Solutions, Inc. (NRSI) and SLR Consulting. Sampling was conducted over a 12-hour period on September 14, 2019, along a 40-meter transect at five sites spanning 1.3 kilometers in clear, free-flowing water. eDNA was collected using two commercially available sampling backpacks (eDNA samplers): (1) the Halltech OSMOS and (2) the Smith-Root ANDe. Electrofishing took place following eDNA collection to avoid any sample contamination that might result. Specimen collection via electrofishing took place in a downsteam-to-upstream orientation along the transect. Four biological replicates were taken at each sampling site, with two samples per eDNA sampler. For each biological sample, six technical replicates were taken. In addition to water samples and abundance counts, abiotic characteristics of the sampling sites were collected, such as air and water temperatures, water pH readings, water dissolved oxygen levels, water conductivities, and water volumes. eDNA was isolated from water filters and amplified via Quantitative Polymerase Chain Reaction (qPCR) in a highly controlled laboratory setting to avoid sample contamination and mitigate qPCR inhibition, among other concerns. Further methodological details, including calculation of eDNA concentrations from standard curve measurements, can be found in Nolan et al. (2023).

Both electrofishing and eDNA sampling revealed the presence of brook trout in Hanlon Creek (Nolan et al., 2023). eDNA was present at all five sampling sites. However, all but Site 1 produced successful electrofishing events (between 1-10 individuals caught; *n* = 15 fish).

#### 2.2.2 Data Discretization

The variables used for rule mining, selected from the Nolan et al. (2022) data set, are shown in **Figure 1** and **Table 1**. Association rules require that continuous attributes be discretized prior to mining, which was accomplished by binning observations into discrete binary categories (“low” and “high”) using biologically meaningful thresholds.

**Figure 1.**
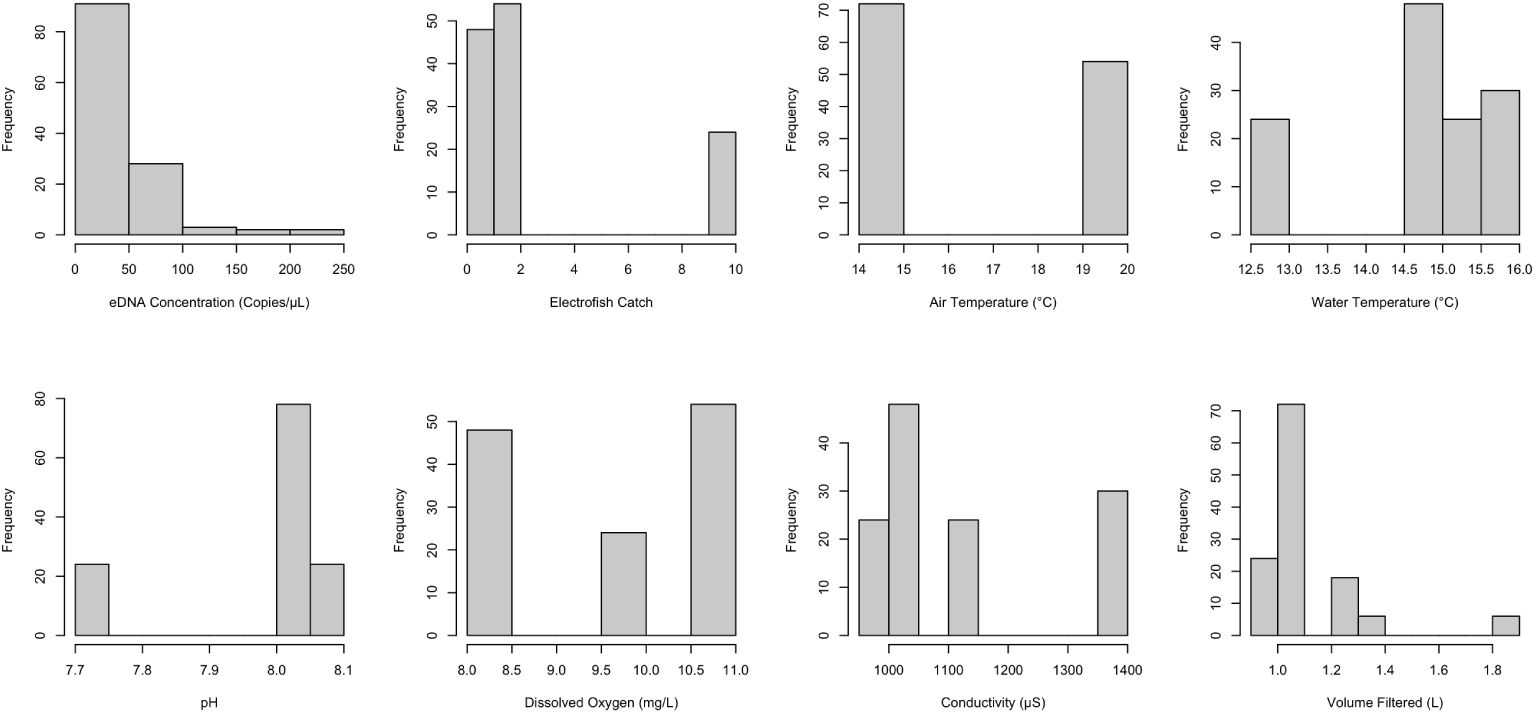
Distributions of quantitative variables used for association rule generation. The data show pronounced skewness across several variables.

**Table 1.**
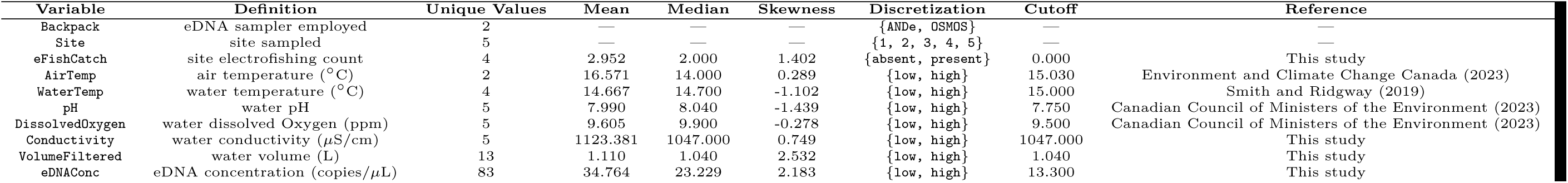
Variable summary statistics (rounded to three decimal places) used to inform association rule mining of brook trout metadata. Negative values of skewness indicate data are negatively (left) skewed, whereas positive values indicate observations are positively (right) skewed.

Determining appropriate thresholds for discretization is a critical but challenging step to data mining, as these thresholds influence the rules generated from the data. For example, unbalanced classes can lead to biased rule generation by disproportionately inflating or limiting the support of certain observations. There are no standardized guidelines for discretization in data mining, thus threshold selection is somewhat subjective, but domain-specific knowledge can provide sufficient justification for chosen cutoffs. This analysis aimed to use discretization thresholds from literature that were both statistically sound and biologically meaningful.

The Canadian Council of Ministers of the Environment (CCME) publishes water quality guidelines to protect aquatic ecosystems (Canadian Council of Ministers of the Environment, 2023). Thresholds for pH and dissolved oxygen were derived from the CCME’s long-term freshwater guidelines. For pH, the CCME recommends a long-term freshwater range of 6.5 to 9.0, thus the midpoint (7.75) was selected as the cutoff to ensure a balanced division between lower and higher values within this acceptable range. For coldwater species like brook trout, the recommended dissolved oxygen range varies from 6.5 mg/L to 9.5 mg/L depending on the life stage. Although the midpoint of this range (8.0 mg/L) was initially selected as the cutoff, this resulted in nearly all dissolved oxygen concentrations being classified as low. To create more balanced bins, the upper bound of 9.5 mg/L was instead selected. An air temperature threshold of 15.03°C was determined by calculating the daily mean temperature in Guelph for September 2019 using data from Environment and Climate Change Canada (Environment and Climate Change Canada, 2023). This was done to increase overall representativeness by smoothing out data to reduce noise inherent in temperature measurements taken on a single day, *i.e.*, the average temperature on September 14. The water temperature threshold of 15.0°C was chosen from a study on the optimal temperature range for brook trout by Smith and Ridgway (2019). Electrofishing catch was discretized into presence and absence categories, with one or more brook trout being classified as “present” and zero being categorized as “absent.”

The eDNA discretization threshold was initially set at 133 copies/µL based on the limit of detection (LOD) established in Nolan et al. (2023). This value represents the lowest standard sample concentration detected with 95% confidence (Klymus et al., 2020). However, this cutoff resulted in no rules being found for `{eDNAConc=low}`. Further, of the 40 rules successfully mined for `{eDNAConc=high}`, the top 10 rules all had very low support of 3.175% and low confidence of 33.333%, along with an extremely high lift of 8.400, a common artifact of low support. Nolan et al. (2023) suggested that a less stringent LOD may be more appropriate for low-concentration eDNA samples, defining it as the lowest standard sample concentration with a 50% detection rate: 13.3 copies/µL in their study. This lower threshold was ultimately adopted for discretizing the eDNA data.

All remaining variables were discretized around the median due to the skewness of the data (see **Figure 1** and **Table 1**). Since the thresholds presented here were derived from multiple sources, they inherently have varying levels of precision, depending on the methods and criteria used in each study.

**Figure 2** depicts an item frequency plot based on the discretization scheme employed herein. Across all transactions, the antecedent `{pH=high}` and the consequent `{eFishCatch=present}` each occurred with the highest frequency of 80.952% (102/126), whereas `{pH=low}`, Sites 1-4, and `{eFishCatch=absent}` all occurred with the lowest frequency of 19.048% (24/126).

**Figure 2.**
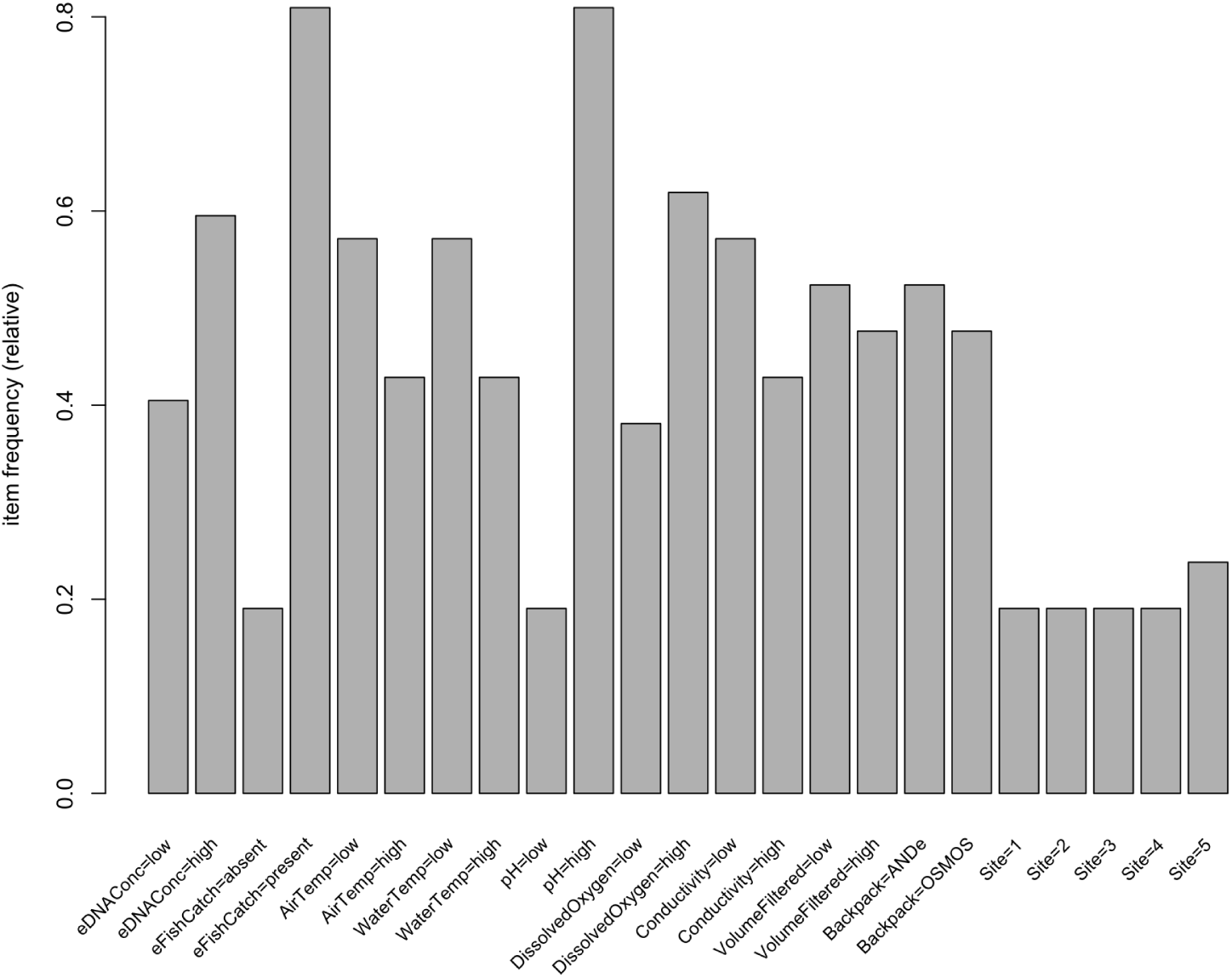
Relative item frequency plot of discretized brook trout environmental covariates.

#### 2.2.3 The RulesTools R Package

To facilitate the rule mining task, we developed the `RulesTools` an R package called (Toth and Phillips, 2025). A link to the package is included in the Data Availability statement. `RulesTools` builds on the functionality of the `arules` R package (Hahsler et al., 2023, 2011, 2005) with additional preprocessing, rule analysis, and result visualization tools. Key functions are:

- `dtize_col()`: Discretizes a numeric column into categorical bins according to user-specified thresholds with native support for missing values (NAs).
- `dtize_df()`: Discretizes a dataframe into categorical values based on specified cutoff points with native support for NAs.
- `compare_rules()`: Compares multiple rulesets to identify association rules common to all possible input set combinations (*e.g.*, rules common to sets A, B, and C; rules common to A and B but not C; rules unique to each set).
- `rule_euler()`: Generates an Euler diagram to illustrate overlaps among multiple rulesets. Note that an Euler plot is similar to a Venn diagram but displays only actual intersections. The degree of overlap indicates the number of elements shared among rulesets.
- `rule_heatmap()`: Generates a heatmap to visualize the association strength between itemsets using support, confidence, or lift as the metric.

Missing values are handled through the `mice` R package, which performs multiple imputation via chained equations (van Buuren and Groothuis-Oudshoorn, 2011). By default, `mice` fits a linear regression model using predictive mean matching (PMM), which uses nearby similar data points to replace missing observations. With respect to eDNA stochasticity, missing values in compiled datasets are presumed to be missing completely at random (MCAR). MCAR assumes that the reason particular data observations are missing is entirely unrelated to both observed and unobserved measurements linked to the dataset. In addition to `mice`, `RulesTools` also relies on well-known R packages in the Tidyverse (Wickham et al., 2019), a suite of R packages for exploratory data analysis, in particular `ggplot2` (Wickham, 2016) for graphics creation. Moreover, `eulerr` (Larsson and Gustafsson, 2018; Larsson, 2024) was used for the generation of Euler diagrams.

The `RulesTools` package greatly facilitated association rule mining throughout this study. Initially, three different discretization schemes were considered for continuous variables: (1) using the mean, (2) using the median, and (3) applying biologically-relevant thresholds derived from primary literature. `dtize_df()` was used to perform these discretizations, and

`compare_rules()` enabled comparisons between the resulting rulesets, identifying rules common across schemes and those unique to each. These comparisons were crucial in guiding the selection of biologically informed thresholds as the most appropriate discretization for the analysis. Furthermore, visualization tools from the package assisted in interpreting rules. `rule_euler()` illustrated how the number of rules progressively decreased through redundancy and significance filtering (see **Figure 3**; see also **Supplementary Files 5-7**). Additionally, `rule_heatmap()` was used to visualize support, confidence, and lift across rules for each of the four consequents, ranking antecedent strength within each metric (see **Figure 4**; see also **Supplementary Files 8-16**).

**Figure 3.**
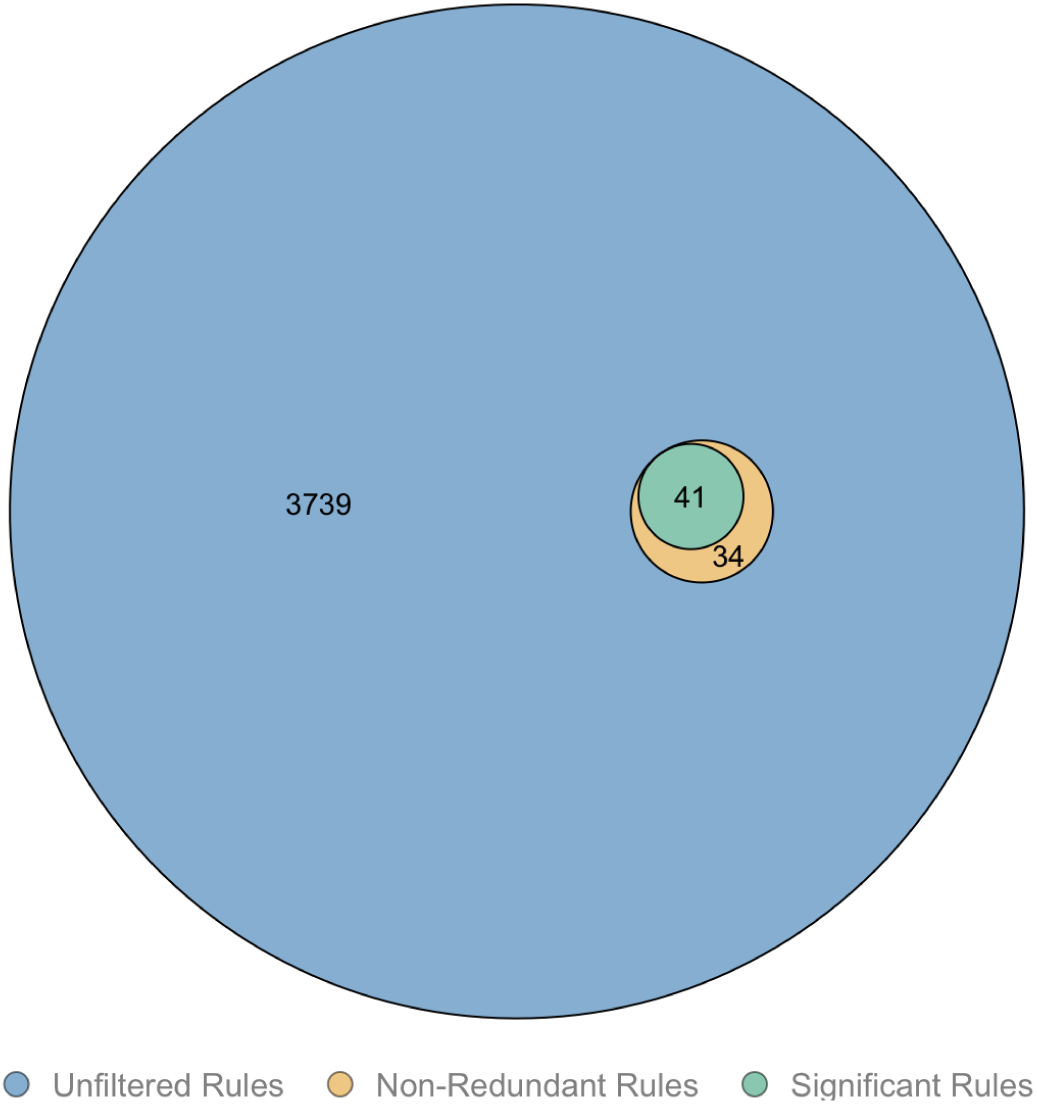
Euler diagram illustrating the pruning of association rules predicting `{eDNAConc=high}`. Set A (blue + yellow + green) represents the initial 3,814 (3739 + 34 + 41) rules generated by the Apriori algorithm. Redundancy filtering reduced this set to 75 non-redundant rules (Set B, yellow + green). Further significance testing at α = 0.05 reduced the set to 41 statistically significant rules (Set C, green). Similar diagrams were generated for the remaining consequents in this study and can be found in **Supplementary Files 5-7**.

**Figure 4.**
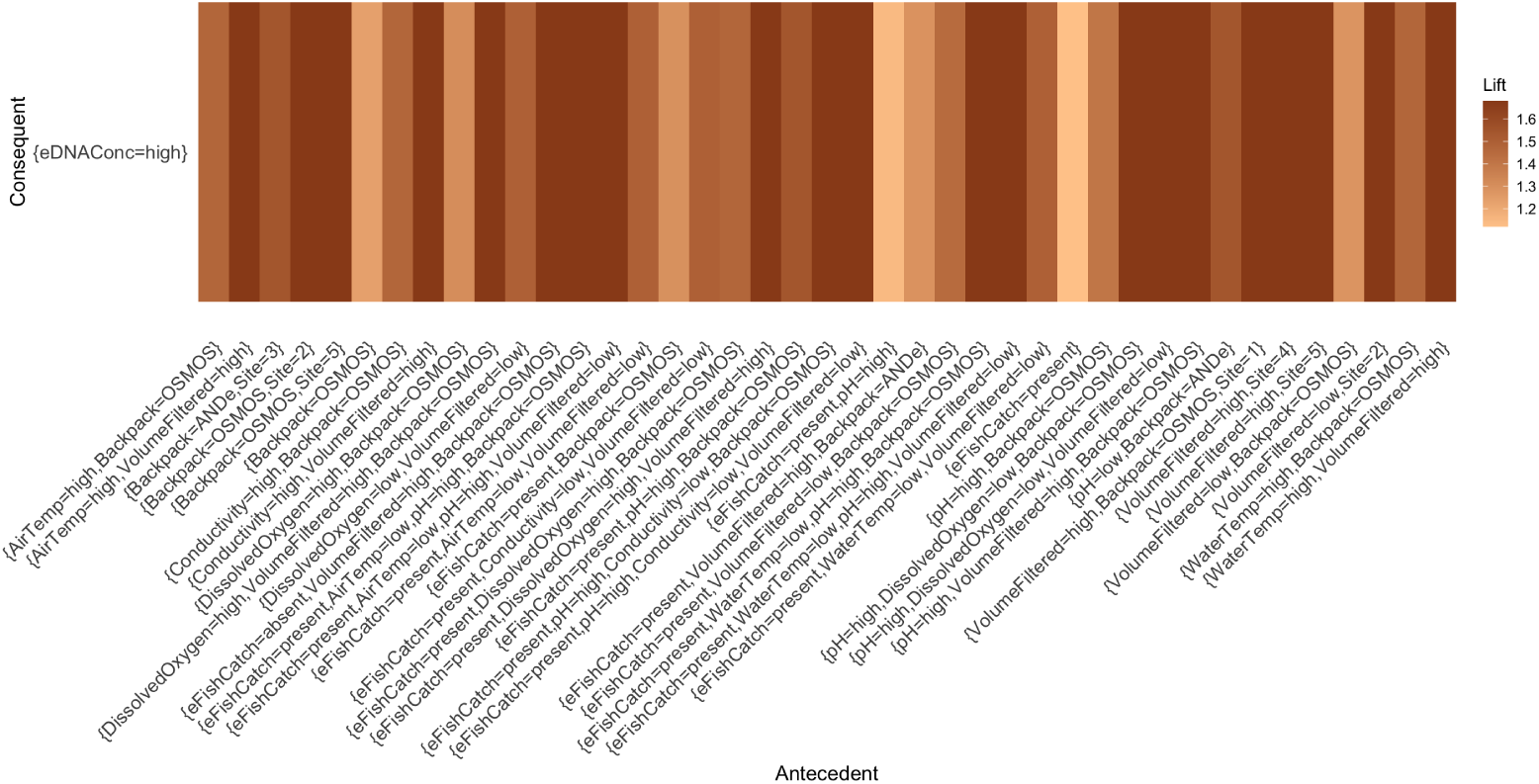
Heatmap depicting the lift of rules generated for the given antecedents and the consequent `{eDNAConc=high}`. Similar diagrams were generated for the remaining rules and interestingness measures in this study and can be found in **Supplementary Files 8-16**.

Future versions of the `RulesTools` package will incorporate associative classification to automate predictive model building, enabling inference of eDNA detection at unsampled sites given particular combinations of abiotic covariates.

#### 2.2.4 Brook Trout Rule Mining

The final dataset comprised 126 transactions and 10 items, with each item represented by two to five levels (23 total levels). Transactions were derived after removing no-template controls, positive controls, internal positive controls and Biomeme qPCR kit results (excluded ue to accuracy concerns, following Nolan et al. (2023)). Note that by equations (1) and (2), this discretization scheme results in 8,388,607 possible itemsets and 94,126,401,612 total association rules. These 126 transactions were derived from four biological replicates per site, each measured with six technical replicates across five sites, plus six additional technical replicates to account for a tear in an ANDe eDNA sampler filter (4 × 6 × 5 + 6 = 126). The `apriori()` function from the `arules` package was used to mine strong association rules from frequent itemsets via the Apriori algorithm (Agrawal and Srikant, 1994). Apriori employs a “bottom-up” approach, progressively constructing more complex rules from simpler ones by first considering all possible 1-itemsets, then 2-itemsets, followed by 3-itemsets, and so forth. The algorithm works under the assumption that the support of an itemset never exceeds the support of its subsets, meaning that if an itemset is frequent, all of its subsets must be frequent as well. Whereas Apriori employs a breadth-first search, other algorithms for frequent item mining—such as ECLAT (Equivalence Class Clustering and bottom-up Lattice Traversal), which is a depth-first search approach—may be more efficient for larger eDNA datasets. All rules were mined with a minimum support of 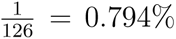 and the same minimum confidence. Note that a threshold of 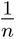, where *n* is the number of database transactions, was selected because it ensures that any rule is supported by at least one transaction in the dataset.

Given the large number of rules generated by the `apriori()` function, all rulesets were further refined through a two-step pruning process to make the data mining task more insightful and manageable: first pruning for redundancy, then testing for significance. In association rule mining, a given rule is considered redundant if it does not provide additional information beyond what is captured by a more general rule with the same or higher value for the chosen interestingness measure (Hahsler and Johnson, 2022). Specifically, a rule *A* ⇒ *C* is redundant if there exists another rule *B* ⇒ *C*, where *B* ⊂ *A* (*i.e.*, *B* is a subset of *A*), and both rules have the same or higher value for the chosen measure. Simplifying rulesets by removing redundancies is advantageous because rules with fewer items are easier to interpret. Thus, rules were filtered for redundancy using the `is.redundant()` function from the `arules` package, with redundancy assessed based on confidence, which is the default. To further refine each ruleset, the `is.significant()` function was applied to test the statistical significance of each rule at a significance level of α = 0.05 (Hahsler and Johnson, 2022). This function evaluates each rule using Fisher’s exact test, which assesses whether the occurrence of the rule’s antecedent and consequent together differs significantly from what would be expected by chance. By default, `is.significant()` makes no adjustment for multiple comparisons, which artificially inflates the Type I error rate, meaning that more rules are retained than should be whenever the null hypothesis of no association between the antecedent and consequent holds. Multiple comparison adjustments were not applied in this study; however, we previously investigated the effect of p-value adjustments such as the Bonferroni method on mined rules for eDNA datasets. (Toth et al., 2026).

Whenever possible, lift was the interestingness measure used to rank resulting association rules, with higher values being favoured. However, its use is not always justified. When multiple rules share identical lifts, rules should be ranked based on confidence. In such cases where mined rules share the same confidence value, support should be employed to break ties.

Following the automated ranking process, rules were manually reviewed to identify any odd, spurious, or biologically implausible associations in need of further scrutiny by domain-specific experts. This step ensured that the resulting rules were not only statistically significant but also ecologically meaningful. Unexpected trends were carefully examined, with particular attention given to associations that deviated from established biological knowledge. In some cases, these anomalies provided novel insights, warranting further investigation, while in others, they indicated potential data artifacts or biases in the discretization process.

## 3 Results

Across all four consequents of interest in this study, a total of 12783 association rules were successfully mined and filtered down to 153 rules. This included rules of the form {} ⇒ *C*, which indicate that the consequent occurs frequently in a given dataset, regardless of the items appearing in the antecedent. As a result, the support and confidence of such rules will be equal. However, these associations are often of little interest; thus, they are usually filtered out.

### 3.1 Rules Predicting Brook Trout Presence/Absence Based on eDNA Concentrations

#### 3.1.1 High eDNA Concentrations

The Apriori algorithm initially generated 3814 rules with the consequent `{eDNAConc=high}`. Redundant rules were removed from the initial ruleset by applying the `is.redundant()` function, reducing the number of rules from 3814 to 75. Finally, the `is.significant()` function was applied at a significance level of α = 0.05, resulting in a final set of 41 rules (**Figure 3**). All rules mined for `{eDNAConc=high}` can be found in **Supplementary File 1**.

Within the set of significant rules, rule lengths ranged from two to five items, with a plurality (19/41; 46.341%) consisting of three items. Rules were supported by between six (6/126 = 0.048) and 68 (68/126 = 0.540) transactions, with a median of 12 (12/126 = 0.095) and a mean of 19.200 transactions. Confidence values, ranged from 0.667 to 1.000, with a median confidence of 1.000 and a mean of 0.919. In other words, there was a 66.700% to 100.000% probability of each rule being true, demonstrating a strong ability of the items in the antecedent to predict high eDNA concentrations. Lift values were between 1.120 and 1.680, with a median lift of 1.680 and a mean of 1.543, reflecting that the items in the antecedent increased the likelihood of high eDNA concentrations by up to 68.000% compared to a baseline of statistical independence (**Figure 4**).

**Table 2** presents the top 10 non-redundant and statistically significant association rules with `{eDNAConc=high}` as the consequent. All rules had a confidence of 1.000, meaning that whenever the antecedent conditions were met, high eDNA concentrations were always observed. The rules shared a support of 0.095, indicating that the items in these rules were observed together in approximately one tenth of all transactions. Additionally, all rules had a lift of 1.680, implying that the antecedents and the consequents were not statistically independent but were instead positively associated. Since all rules share identical confidence, support, and lift values, their ranking in the table is arbitrary and does not imply relative importance or strength.

**Table 2.**
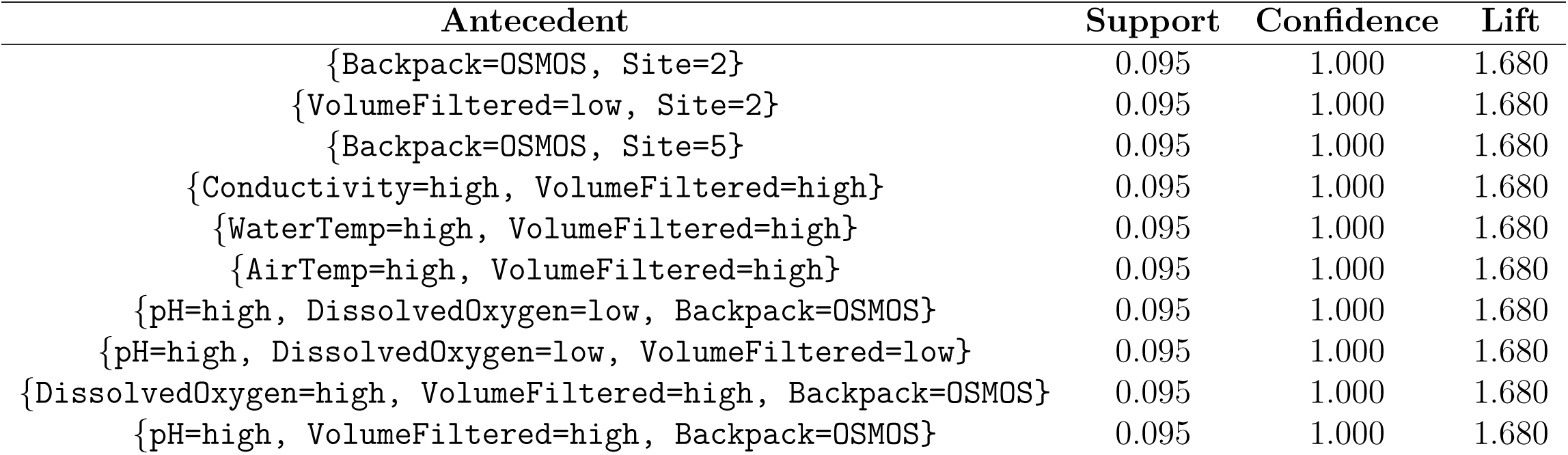
Top 10 mined rules (non-empty, non-redundant, and statistically significant) with the consequent {eDNAConc=high} ranked by lift.

Notable antecedent patterns included the frequent observation of the OSMOS eDNA sampler, which appeared as a subset in half of the top 10 rules (50.000%), suggesting that OSMOS may offer an advantage over the ANDe eDNA sampler in detecting high eDNA concentrations. The condition `{VolumeFiltered=high}` occurred in five of the top 10 rules (50.000%), consistent with the established notion that larger sample volumes generally correspond to higher eDNA concentrations. However, `{VolumeFiltered=low}` also occurred in two of the top 10 rules (20.000%). These associations could be spurious, arising due to the relatively small size of the brook trout dataset used to generate the rules. Despite this, all of these findings support those of Nolan et al. (2023).

#### 3.1.2 Low eDNA Concentrations

The Apriori algorithm generated a preliminary set of 3213 rules for the consequent `{eDNAConc=low}`. Redundant rules were removed, reducing the ruleset to a size of 60. Eliminating non-significant rules using a significance level of α = 0.05 resulted in a final set of 35 significant rules. The complete set of rules mined for `{eDNAConc=low}` can be found in **Supplementary File 2**.

Within this set of 35 rules, the rule lengths ranged from two to five items, with the majority of rules (29/35; 82.857%) consisting of three or four items. The rules were supported by between five (5/126 = 0.040) and 35 (35/126 = 0.278) transactions, with a median of 16 (16/126 = 0.127) and a mean of 14.700 transactions. Confidence values ranged from 0.521 to 1.000 with a median confidence of 0.750 and a mean of 0.785, representing a 52.100% to 100.000% probability of each rule being true. Lift values varied from 1.287 to 2.471, with a median lift of 1.853 and a mean of 1.940, reflecting that the antecedent conditions were up to 147.100% more likely to occur alongside low eDNA concentrations compared to a baseline of statistical independence.

The top 10 non-redundant, significant rules with the consequent `{eDNAConc=low}` are shown in **Table 3**. These rules had support values ranging from 0.048 to 0.095, indicating the associations were observed in approximately one out of every 10 to 20 transactions. All the confidence values were 1.000, suggesting perfect reliability. All of the lift values were 2.471, indicating that the antecedent conditions significantly increased the likelihood of low eDNA concentrations compared to a condition of statistical independence.

**Table 3.**
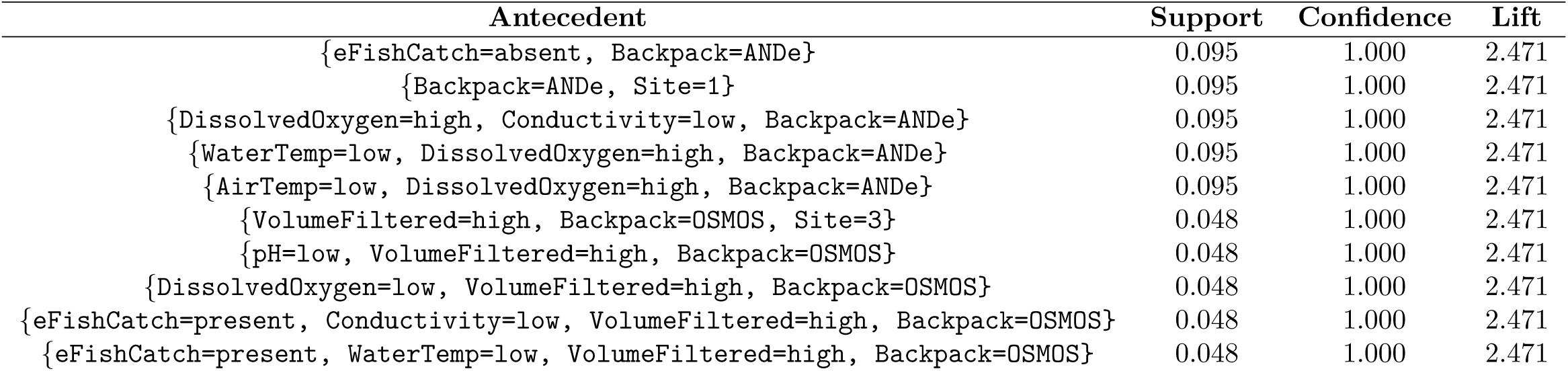
Top 10 mined rules (non-empty, non-redundant, and statistically significant) with the consequent {eDNAConc=low} ranked by lift.

Several key antecedent patterns emerged from these top 10 rules, revealing a complex and somewhat contradictory relationship with low eDNA concentrations. Unlike the rules for high eDNA concentrations, which consistently showed that the OSMOS sampler yielded higher levels of eDNA, the sampler type in these rules was evenly split: ANDe appeared in five rules (50.000%), and OSMOS appeared in five rules (50.000%). This suggests that for low eDNA concentrations, the influence of the sampler type is less clear-cut. A similar contradiction arose with the `eFishCatch` variable. In these rules, `eFishCatch` appeared as present in two rules (20.000%) but as absent in another rule (10.000%). This variability may reflect the limited size of the dataset. Nevertheless, some patterns aligned with previous findings. For example, the rule:

`{Backpack=ANDe, Site=1} ⇒ {eDNAConc=low}`

corroborated findings from Nolan et al. (2023) where electrofishing at Site 1 yielded no brook trout. Furthermore, the rule

`{eFishCatch=absent, Backpack=ANDe} ⇒ {eDNAConc=low}`

was intuitive, as low electrofishing yields generally suggest brook trout absence and therefore lower eDNA concentrations.

### 3.2 Rules Predicting Brook Trout Presence/Absence via Electrofishing

#### 3.2.1 Brook Trout Presence

Mining targeted the consequent `{eFishCatch=present}` to identify association rules predicting successful electrofishing events, providing a point of comparison with eDNA sampling. A total of 4412 rules were generated by the Apriori algorithm. Removing redundant rules reduced this to 17 rules, and further statistical significance testing at α = 0.05 resulted in a final set of 14 rules. All rules mined for `{eFishCatch=present}` can be found in **Supplementary File 3**.

The rules exhibited lengths between two and three items, with the majority (11/14; 78.571%) consisting of two items. The rules were supported by between 24 (24/126 = 0.191) and 68 (68/126 = 0.540) transactions, with a median of 35 (35/126 = 0.278) and a mean of 39 transactions. Confidence values ranged from 0.907 to 1.000, with a median of 1.000 and a mean of 0.985, suggesting high reliability in predicting brook trout presence. The lift values spanned from 1.120 to 1.235, with a median lift of 1.235 and a mean of 1.217, highlighting that antecedent conditions were 12.000% to 23.500% more likely to occur alongside the consequent `{eFishCatch=present}` compared to a baseline of statistical independence.

**Table 4** shows the top 10 non-redundant and statistically significant association rules with `{eFishCatch=present}` as the consequent. All rules were found to have a confidence of 1.000, indicating that whenever the antecedent conditions were met, brook trout presence was detected via electrofishing. The support values for these rules ranged from 0.190 to 0.429, suggesting that these associations appeared in a substantial subset of transactions. The lift values were each 1.235 meaning that the rules slightly increased the likelihood of successful electrofishing outcomes compared to what would be expected if the rule items were statistically independent.

**Table 4.**
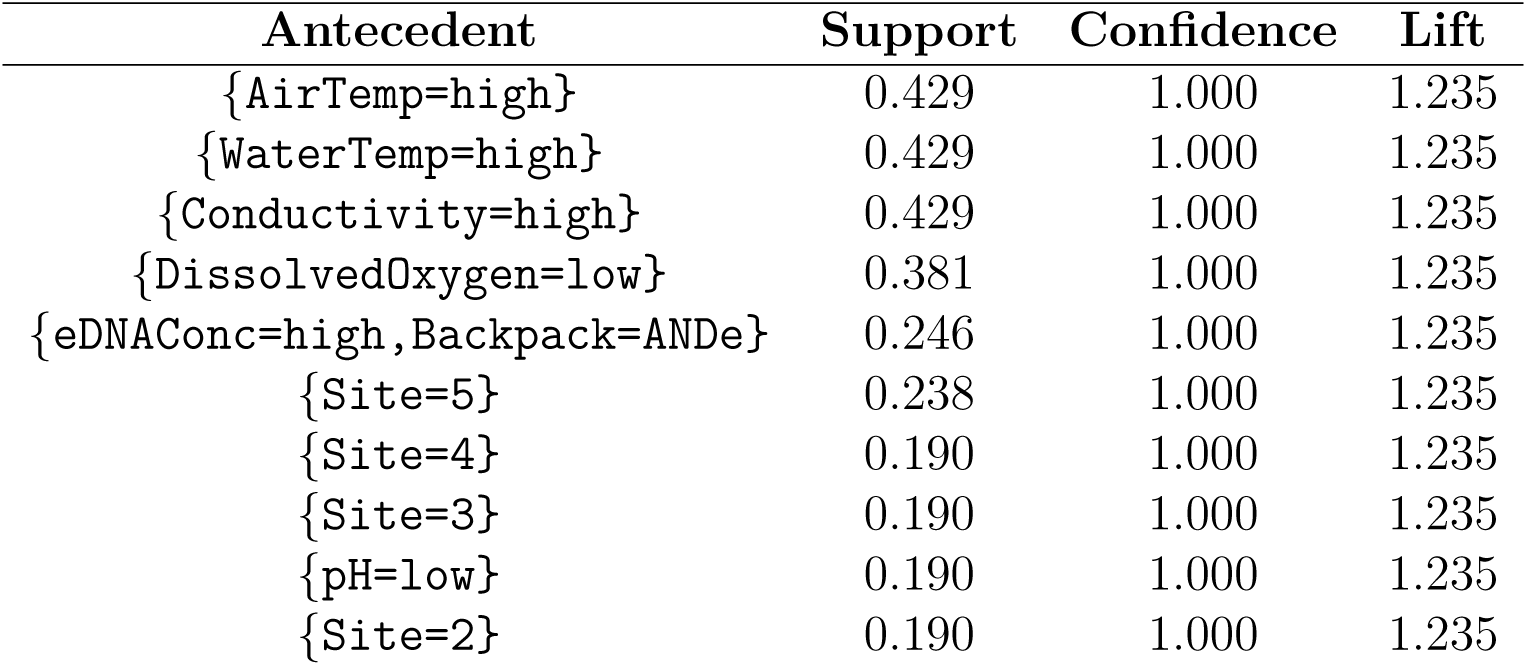
Top 10 mined rules (non-empty, non-redundant, and statistically significant) with the consequent {eFishCatch=present} ranked by lift.

It is notable that setting the consequent to `{eFishCatch=present}` generated simpler rules with no more than two items in the antecedent. Interestingly, there also did not appear to be a consistent relationship between brook trout presence detected by electrofishing and eDNA concentrations. Only one rule in the top 10 (10.000%) featured `{eDNAConc=high}` in the antecedent. Nolan et al. (2023) generally found correspondence between brook trout caught via electrofishing and those detected through eDNA, with brook trout netted at four of the five (80.000%) sites within Hanlon Creek and eDNA detected at all five sites (100.000%).

#### 3.2.2 Brook Trout Absence

The final step of this analysis was mining association rules predicting the absence of brook trout via the consequent `{eFishCatch=absent}`. Initially 1344 rules were mined. This ruleset was reduced to 67 rules by removing redundancies, and further reduced to a set of 63 rules by removing statistically non-significant rules. All rules with the consequent `{eFishCatch=absent}` can be found in **Supplementary File 4**.

Rule lengths within this set ranged between two and five items; the majority (43/63; 68.254%) consisted of three or four items. The rules were supported by between five (5/126 = 0.040) and 24 (24/126 = 0.190) transactions, with a median of 12 (12/126 = 0.095) and a mean of 13.500 transactions. The confidence values varied between 0.235 and 1.000, with a median of 0.600 and a mean of 0.647, showing a 23.500% to 100.000% chance of each rule being true. The lift values ranged from 1.235 to 5.250, having a median of 3.150 and a mean of 3.394, revealing strong positive associations between the antecedents and brook trout absence.

**Table 5** shows the top 10 non-redundant and statistically significant association rules with `{eFishCatch=absent}` as the consequent. All rules were found to have a confidence of 1.000, meaning that whenever the antecedent conditions were met, brook trout absence was consistently observed. The support values for these rules ranged from 0.048 to 0.190. The lift values were each 5.250, reflecting that the antecedent conditions increased the likelihood of brook trout absence by 425.000% compared to random chance.

**Table 5.**
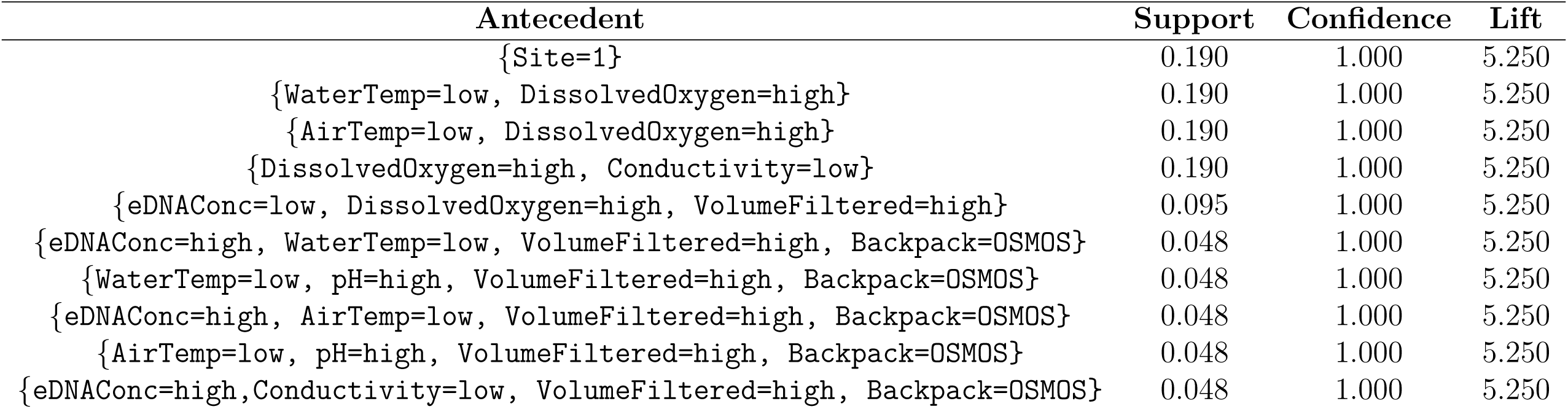
Top 10 mined rules (non-empty, non-redundant, and statistically significant) with the consequent {eFishCatch=absent} ranked by lift.

The top rule featured Site 1 as the antecedent, which was expected since Site 1 was the only site where no brook trout were caught (Nolan et al., 2023). However, there were also three rules where high eDNA concentrations appeared in the antecedent. The presence of high eDNA concentrations may once again be spurious due to the limited dataset size.

## 4 Discussion

This study demonstrates the utility of association rule mining for uncovering interesting relationships among environmental metadata variables from eDNA sampling and electrofishing datasets. Using data collected from Hanlon Creek in Guelph, ON, associations linking environmental covariates and eDNA sampling methods to brook trout detections were mined. The resulting rules can be used to predict sampling outcomes and identify the conditions most conducive to successful data collection.

### 4.1 Analysis of Rules Predicting High Brook Trout eDNA Concentrations

Rules mined with the consequent `{eDNAConc=high}` revealed several interesting associations, some of which were consistent with existing eDNA literature, while others contradicted expectations. Of the 41 rules generated, only one (1/41 = 2.439%) featured `{pH=low}` in the antecedent while 12 (12/41 = 29.268%) included the item `{pH=high}`. This pattern aligns with existing research showing that eDNA is more persistent in alkaline environments, where lower rates of acid hydrolysis and slower enzymatic activity delay degradation compared to acidic environments (Seymour et al., 2018).

Water temperature’s role in the rules was more nuanced than existing literature would suggest. Although lower temperatures have consistently been shown to preserve eDNA due to reduced microbial and chemical activity (McCartin et al., 2022), water temperature was only featured in the antecedents of five (5/41 = 12.195%) rules. Two (2/41 = 4.878%) of these linked high eDNA concentrations with *high* temperatures. Several past studies have noted that elevated temperatures promote greater organism eDNA shedding rates across taxa (*e.g.*, Andruszkiewicz Allan et al. (2021); Harrison et al. (2019); Kirtane et al. (2021); Sassoubre 8 et al. (2016)). These findings highlight the necessity of applying domain-specific knowledge when interpreting association rules as well as when setting appropriate minimum thresholds for data mining. Although data mining can reveal unexpected patterns, results that contradict established knowledge may reflect spurious associations due to small sample sizes, stochasticity, or unidentified confounding variables. In this study, where the dataset comprised just 126 transactions, it is likely that these counterintuitive rules were mere chance associations despite the stringent pruning system used.

Overall, rules produced for the consequent `{eDNAConc=high}` scored well across all evaluation metrics. Confidence was never lower than 0.667, with a median of 1.000 and a mean of 0.919, indicating that the antecedents were strongly and consistently linked to high eDNA concentrations. The lowest lift value of 1.120 indicated that all antecedents increased the likelihood of high eDNA concentrations by at least 12.000% compared to random chance. Although several rules had low support, their high confidence and lift suggest that these associations, though infrequent, still reliably indicated conditions conducive to high eDNA concentrations.

### 4.2 Analysis of Rules Predicting Low Brook Trout eDNA Concentrations

Rules with the consequent `{eDNAConc=low}` were weaker and less consistent with established literature compared to those mined for the consequent `{eDNAConc=high}.` For instance, higher water temperatures are known to promote greater eDNA degradation (McCartin et al., 2022; Tsuji et al., 2017). However, seven of the 35 final rules (7/35 = 20.000%) linked low eDNA concentrations with low water temperatures, while the `antecedent {WaterTemp=high}` did not appear at all. Furthermore, seven rules (7/35 = 20.000%) featured `{pH=high}` in the antecedent while only one (1/35 = 2.857%) contained the item `{pH=low},` contradicting the established understanding that acidic conditions promote greater eDNA degradation (Seymour et al., 2018). Several of the rules that included `{pH=high}` or `{WaterTemp=low}` also involved additional covariates. For example, the rules

`{pH=high, Conductivity=low, Backpack=ANDe} ⇒ {eDNAConc=high}`

and

`{WaterTemp=low, DissolvedOxygen=high, Backpack=ANDe} ⇒ {eDNAConc=high}`

featured conductivity, dissolved oxygen, and eDNA sampler type. These multi-variable rules may obscure the individual influences of pH or water temperature on eDNA concentrations and may instead imply that complex interactions among environmental factors dictate sampling outcomes. Such patterns highlight the importance of considering items in association rules in the context of the full antecedent.

Rules predicting low eDNA concentrations performed well across key interestingness metrics, though they were slightly weaker than those predicting high eDNA concentrations. Median and mean confidence were 0.750 and 0.785, respectively—slightly lower than the near-perfect confidence of the `{eDNAConc=high}` rules, but still indicative of strong and reliable associations. Support was also slightly lower for `{eDNAConc=low}` rules, with a range of 0.040 to 0.278, suggesting that these patterns occurred less frequently but were still meaningful within the dataset. Lift was the only measure for which the low concentration rules performed better than the high concentration rules, with values up to 2.471 indicating that associations were up to 147.100% more likely to occur than by random chance.

Interestingly, several rules associated low eDNA concentrations with low conductivity, but the antecedent `{Conductivity=high}` did not appear in any of the 35 rules. Although the effect of water conductivity on eDNA persistence is not well characterized, a study by Collins et al. (2018) showed a negative correlation in marine environments, which would potentially support the rules generated in this study. Collins et al. (2018) suggest that low conductivity conditions may indirectly affect eDNA concentrations by supporting microbial communities that facilitate degradation. Notably, the rule

`{Conductivity=high, VolumeFiltered=high} ⇒ {eDNAConc=high}`

from **Table 2** suggests the same trend, by conversely linking high water conductivity with high eDNA concentrations. Nevertheless, literature examining the impact of conductivity on eDNA concentrations in freshwater ecosystems is sparse, and the results of this data mining effort suggest that the relationship between conductivity and eDNA persistence warrants further investigation.

Additionally, the rules

`{eFishCatch=present, Conductivity=low, VolumeFiltered=high, Backpack=OSMOS} ⇒ {eDNAConc=low}`

and

`{eFishCatch=present, WaterTemp=low, VolumeFiltered=high, Backpack=OSMOS} ⇒ {eDNAConc=low}`

suggest false negative detection, indicating that developed molecular assays may lack sufficient sensitivity and specificity that enable the robust detection of aquatic eDNA signal.

### 4.3 Analysis of Rules Predicting Brook Trout Presence via Electrofishing

One of the most interesting trends that emerged among the rules predicting `{eFishCatch=present}` was the simplicity of the final ruleset. Only 14 rules remained after pruning and the majority of them (11/14 = 78.571%) featured a single item in the antecedent. These concise rules may reflect a class imbalance introduced by discretizing electrofishing data into presence and absence categories, where `{eFishCatch=present}` occurred 102 times (102/126 = 80.952%) and `{eFishCatch=absent}` occurred only 24 times (24/126 = 19.048%). Because brook trout were present in most transactions, many simple, single-variable conditions were sufficient to meet minimum support and confidence thresholds. As a result, the ruleset might favour patterns associated with this majority class. The bias toward over-represented classes in association rule mining presents a challenge for interpreting results, as it can lead to over-confidence in simpler rules while masking meaningful patterns associated with minority classes. To mitigate this, it is often necessary to consider class balancing techniques or alternative interestingness measures. For instance, rules could be ranked according to measures designed to handle unbalanced classes such as local support, which measures the proportion of transactions containing the consequent where the rule’s antecedent conditions also hold (Gu et al., 2003).

### 4.4 Analysis of Rules Predicting Brook Trout Absence via Electrofishing

After pruning, a total of 63 rules predicting `{eFishCatch=absent}` remained. Some of these associations were likely spurious. For instance, 24 rules (24/63 = 38.095%) included Backpack as an antecedent item, but these rules were deemed irrelevant because there was no plausible mechanism by which eDNA sampler type would influence electrofishing results. Other associations were more meaningful. For example, 29 rules (29/63 = 46.032%) featured `{eDNAConc=low}`, while only four (4/63 = 6.349%) included `{eDNAConc=high}`, which revealed an expected and logical positive correlation between electrofishing catch and eDNA concentrations.

The average confidence for `{eFishCatch=absent}` rules was lower than for any other consequent, with a median of 0.600 and a mean of 0.647. Because unsuccessful electrofishing outcomes occurred in just 24 of 126 (19.048%) transactions, it co-occurred with all antecedents somewhat infrequently, lowering overall confidence scores. Support was also low, ranging between 0.040 and 0.240. Conversely, lift was exceptionally high for these rules—upwards of 5.250 which can also be attributed to the low frequency of `{eFishCatch=absent}` in the dataset. Since the support of the antecedent constitutes the denominator of the lift equation, an exceptionally low support can amplify the result. Thus, it is important to caution that while high lift can often indicate strong associations, in imbalanced datasets it may overstate the importance of rules with low support. These findings reflect the challenges associated with predicting minority classes.

The rules

`{eDNAConc=high, WaterTemp=low, VolumeFiltered=high, Backpack=OSMOS} ⇒ {eFishCatch=absent},`

`{eDNAConc=high, AirTemp=low, VolumeFiltered=high, Backpack=OSMOS} ⇒ {eFishCatch=absent},`

and

`{eDNAConc=high, Conductivity=low, VolumeFiltered=high, Backpack=OSMOS} ⇒ {eFishCatch=absent},`

suggest a mechanism related to downstream eDNA transport. As creeks comprise lotic (fast moving) water bodies, eDNA could have originated from further upstream where brook trout were observed through electrofishing. This finding reinforces the need to measure flow rates (which were not collected for the Hanlon Creek data). The last rule above in addition is suggestive that electrofishing was unsuccessful due to low water conductivity. Freshwater environments generally possess lower concentrations of ionic salts, which inhibits the movement of strong electrical currents necessary to stun fish; thus, higher voltages are required in these settings (Kolz, 2006).

### 4.5 Implications of Association Rule Mining for eDNA Datasets

Association rule mining holds immense potential as a novel data analysis technique in eDNA studies because it offers several advantages over traditional statistical methods like regression or correlation analysis.

Firstly, rule mining, and unsupervised machine learning in general, is an exploratory method that can be applied absent of any presuppositions about the data, unlike hypothesis-driven statistical methods (Agrawal et al., 1993). The approach can be viewed as an abductive logical tool (as opposed to an inductive or deductive one) since it aims to identify the most plausible relationships among a set of known inputs that together account for an observed output, even when these inferred relationships are not known with complete certainty. The benefit of this approach is that it promotes the discovery of unexpected associations that may have otherwise been overlooked. It is especially useful for detecting complex, multi-variable relationships. Note that after calculating the Spearman correlation coefficients between key environmental variables and eDNA concentrations in this study (see **Figure 5**), none of the correlations were statistically significant at the 5% level, suggesting that no isolated factor had a strong relationship with eDNA levels. However, association rule mining revealed many interesting context-dependent and multi-dimensional relationships among support, confidence, and lift across the four consequents considered in this study (**Figure 6**). Traditional correlation analysis assumes independence among data observations and experiences difficulties when replicated values are present within datasets. These findings underscore the value of data mining for uncovering complicated associations between eDNA concentrations and environmental variables that standard correlation techniques often struggle to detect.

**Figure 5.**
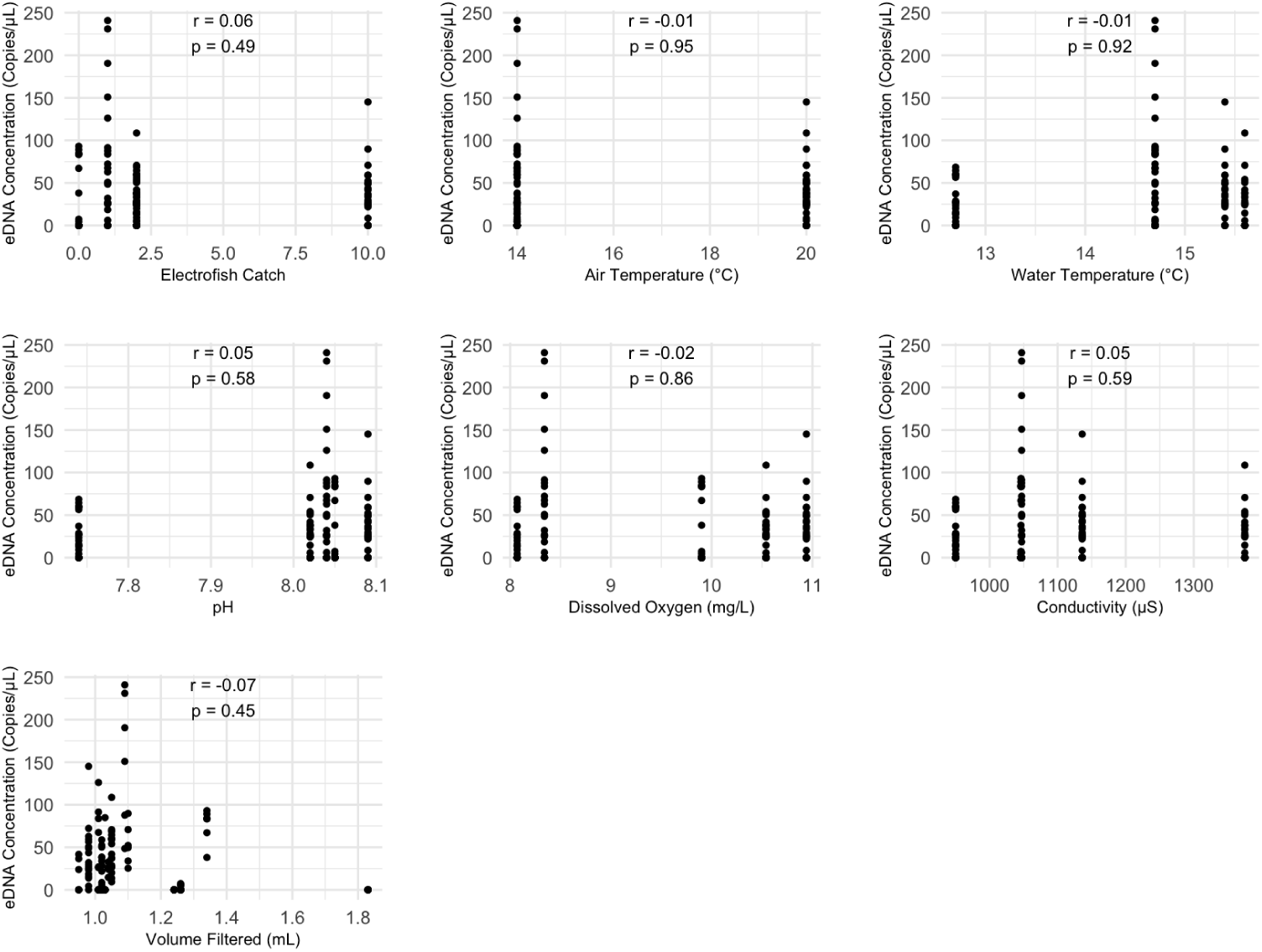
Scatterplots showing the relationship between eDNA concentration (copies/µL) and key environmental variables. Each subplot includes the Spearman correlation coefficient (r) and associated p-value. The discreteness of the data reflects repeated measurements at the same values.

**Figure 6.**
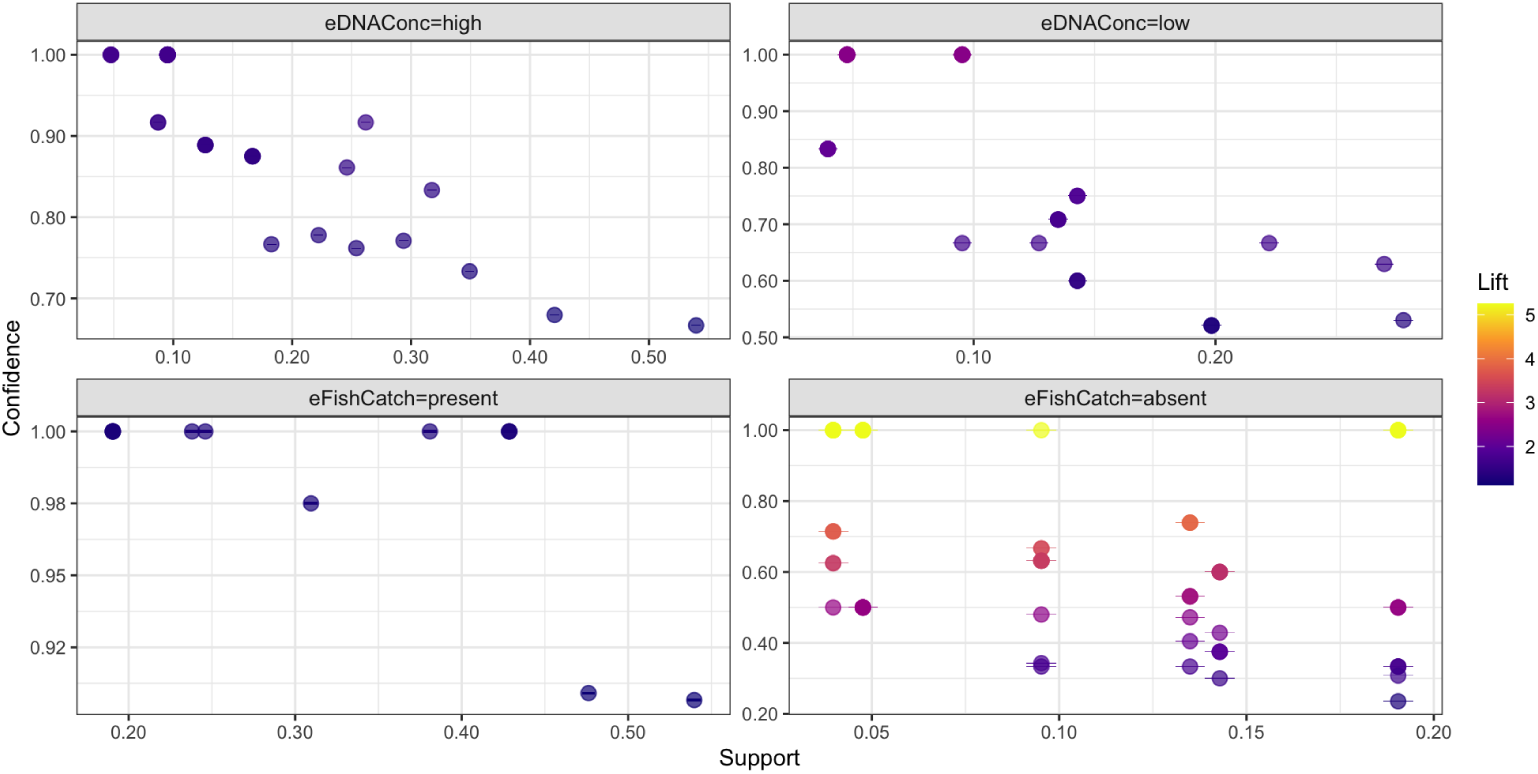
Scatterplots depicting trends across four considered consequencts in this study. Many rules were tied in support, confidence, and lift.

Secondly, association rule mining is blind to underlying data distributions, unlike conventional statistical methods which often require that data follow specific parametric assumptions to generate valid inferences. Association rule mining can be applied to any transactional dataset, even when that dataset’s distribution is unknown or difficult to assess (Han et al., 2011). This is particularly beneficial for the complex and irregular data commonly collected within ecological studies.

Finally, association rules are a highly accessible way to express relationships within datasets. Their simple “if-then” structure makes them easy to interpret even for non-technical audiences (Hahsler et al., 2011). For eDNA research, and environmental research more broadly, this accessibility could support informed decision-making and promote greater public engagement with biodiversity research.

### 4.6 Limitations of Association Rule Mining for eDNA Datasets

While association rule mining offers several indispensable advantages as a data analysis technique, the results of this paper highlight notable limitations that researchers should be aware of when applying the method to eDNA datasets.

First, determining appropriate thresholds for support, confidence, lift, and other interestingness measures is highly subjective and no universal guidelines exist (Tan et al., 2018). Instead, thresholds are chosen based on the particular dataset properties, domain knowledge, and the specific objective of the analysis; however, these choices significantly influence obtained results. For instance, setting thresholds too low can generate a large number of non-significant or trivial rules, while setting cutoffs too high might obscure rare but meaningful associations. Data requiring discretization will introduce additional subjectivity. Alternative approaches to association rule mining, such as fuzzy-based rule systems, which do not require continuous data to first be discretized, offer promising ways forward. Cutoff values and binning strategies alter the structure and frequency of items in the dataset, which significantly impacts which rules are discovered during mining. In this analysis, the imbalance between classes `{eFishCatch=present}` and `{eFishCatch=absent}` resulted in comparatively stronger rules for the former, even though the underlying relationships were not necessarily stronger.

Overfitting is another concern in association rule mining. Rules that capture associations specific to a particular dataset may not generalize well to unseen data, especially if the originally mined dataset is small, noisy, or highly complex (*i.e.*, too many antecedent items) (Ying, 2019). The brook trout dataset used for this analysis was exceptionally small, with just 126 transactions; thus, many of the unexpected relationships uncovered were likely spurious associations resulting from the characteristics of those particular transactions. As association rule mining is extended to other potentially larger datasets, subsequent work will further seek to minimize the presence of redundancies and difficult-to-interpret rules as part of the pruning workflow by mining a concise representation of itemsets. This will entail ensuring that all mined frequent itemsets are both closed and maximal. This can be accomplished via the functions `is.closed()` and `is.maximal()` in `arules`. An itemset is said to be closed if none of its immediate supersets has the same support. Likewise, a maximal itemset is one where none of its immediate supersets is frequent. Thus, all maximal itemsets are also closed. The pruning workflow adopted herein performed reasonably well. For non-redundant and statistically significant rules with the consequent `{eDNAConc=high},` 70.732% (29/41) of generated rules comprised maximal (and hence also closed) itemsets, whereas only 68.571% (24/35) of rules were found to contain maximal itemsets for the consequent `{eDNAConc=low}`. Similarly, 85.714% (12/14) of rules having the consequent `{eFishCatch=present}` were generated from maximal itemsets, while only 39.683% (25/63) of rules with the consequent `{eFishCatch=absent}` contained maximal itemsets. Based on these findings, focus should be placed on retaining only rules formed from maximal itemsets moving forward. However, while association rules generated from closed and maximal itemsets are compact, results may be less generalizable overall. This can be seen in the resulting association rules for electrofishing, where considerable class imbalance was noted. Therefore, caution is necessary, especially when employing mined rules for downstream predictive classification, even though model accuracy is not likely to suffer (Antonie et al., 2016).

Lastly, it is important to highlight that association rule mining is not inherently a statistical method. While it can be used to uncover strong associations, to determine whether or not uncovered relationships are real, further statistical testing may be warranted (as was done in this study by applying Fisher’s exact test to prune non-significant rules). However, because thousands of Fisher’s exact tests were conducted across the four rulesets without correction for multiple comparisons, the expected number of false positives among the 153 retained rules is likely non-trivial, and the reported significance levels should be interpreted as descriptive rather than confirmatory. Small datasets are especially prone to producing spurious rules that do not translate well to unseen data. This further highlights the importance of expert validation in interpreting mined rules (Hahsler et al., 2011), especially since co-occurrence among ruleset items within the antecedent and consequent is not causative. Ascertaining the direct causes of eDNA and species presence/absence would require alternative tools capable of modelling categorical data, such as Bayesian networks.

Despite these limitations, association rule mining can still provide valuable insights, especially when used in combination with other statistical methods such as habitat occupancy modelling to estimate probabilities of species detection and occupancy.

## 5 Conclusion

This paper serves as a proof-of-concept for applying association rule mining to eDNA datasets. Using data collected from Hanlon Creek in Guelph, ON., Canada with 126 transactions across 10 diverse biotic and abiotic environmental covariates, rules were mined to predict brook trout detections via eDNA sampling and electrofishing. The resulting rules performed strongly across key evaluation metrics such as support, confidence, and lift. Furthermore, they revealed interesting, often unintuitive relationships between sampling outcomes and environmental metadata variables, including water conductivity, temperature, and pH, among others. Herein, an LOD of 13.3 copies/µL was used to discretize eDNA concentrations into low and high categories, since a threshold of 133 copies/µL failed to generate any rules with `{eDNAConc=low}` as the consequent. However, an argument could be made for an even tighter LOD threshold of zero, whereby cases of no eDNA detection can be ascertained.

Association rule mining is a versatile unsupervised machine learning technique that holds promise for supporting and improving eDNA data collection and analyses. For instance, rules relating eDNA detections with specific environmental covariates could be used to identify which metadata variables to collect alongside eDNA samples. Poor metadata reporting has been identified as a common weakness among many existing eDNA studies (Nicholson et al., 2020). Additionally, rule mining could be used to complement statistical modelling by highlighting significant covariate combinations to include in regression or occupancy models.

Researchers who wish to use association rule mining for ecological applications should be aware of the challenges and considerations detailed in this paper. These include the subjectivity of threshold selection, the risk of overfitting, and the lack of statistical significance in generated rules. Careful validation of outputted rules is critical, especially for small datasets where spurious associations are likely to arise. Domain-specific expertise should be called upon at every step of the process.

This study lays the groundwork for future applications of machine learning to eDNA datasets, especially in an unsupervised setting. One promising direction is using mined rules to build an associative supervised classifier to enable predictions regarding eDNA concentrations at yet unsampled ecological sites based on collected environmental metadata variables. Overall model performance would then be assessed using well-established evaluation metrics such as accuracy, precision, recall, and *F*_1_-score to mitigate false positive (a species’ DNA being present in a given environmental sample despite said species not occurring at a particular sampling location) and false negative (a species’ DNA not being captured in a given environmental sample even though said species occurs naturally at a given sampling site) detections. Future studies with larger and more diverse datasets, such as those based on eDNA metabarcoding to characterize entire species communities, could also refine and expand the approach developed in this work. Ultimately, association rule mining is an emerging methodology within the ecological literature that can enhance eDNA sampling workflows to support biodiversity research and promote more effective decision-making for conservation and management practices.

## 6 Acknowledgements

We acknowledge that the University of Guelph resides on the ancestral lands of the Attawandaron people and the treaty lands and territory of the Mississaugas of the Credit. We recognize the significance of the Dish with One Spoon Covenant to this land and offer our respect to our Anishinaabe, Haudenosaunee and Métis neighbours as we strive to strengthen our relationships with them.

We would like to thank Kathleen (Kat) Nolan for her generosity, for her invaluable guidance throughout this project, and for providing constructive feedback on a previous draft of this work.

## 7 Author Contributions

NT and JDP wrote the manuscript, wrote required code, as well as analyzed and interpreted all experimental results. LA served as an advisor in data mining and machine learning. RHH acted as an advisor in eDNA sampling. DJG served as an advisor in statistics and data science. All authors contributed to the revision of this manuscript and approved the final version.

