## Supplementary figures and images for "Mining association rules for targeted spatiotemporal aquatic environmental DNA (eDNA) sampling"

### SupplementaryFile5.png

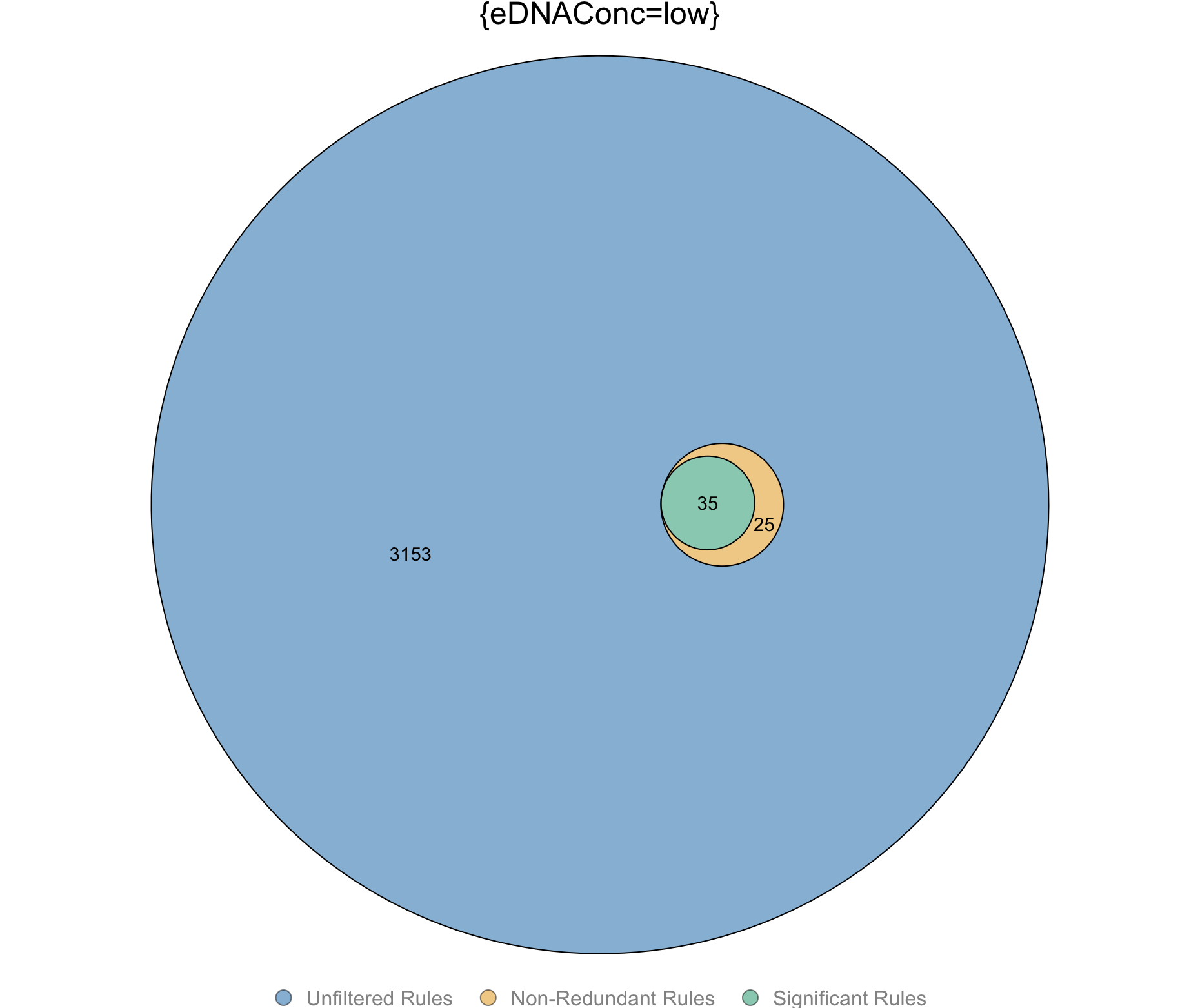

### SupplementaryFile6.png

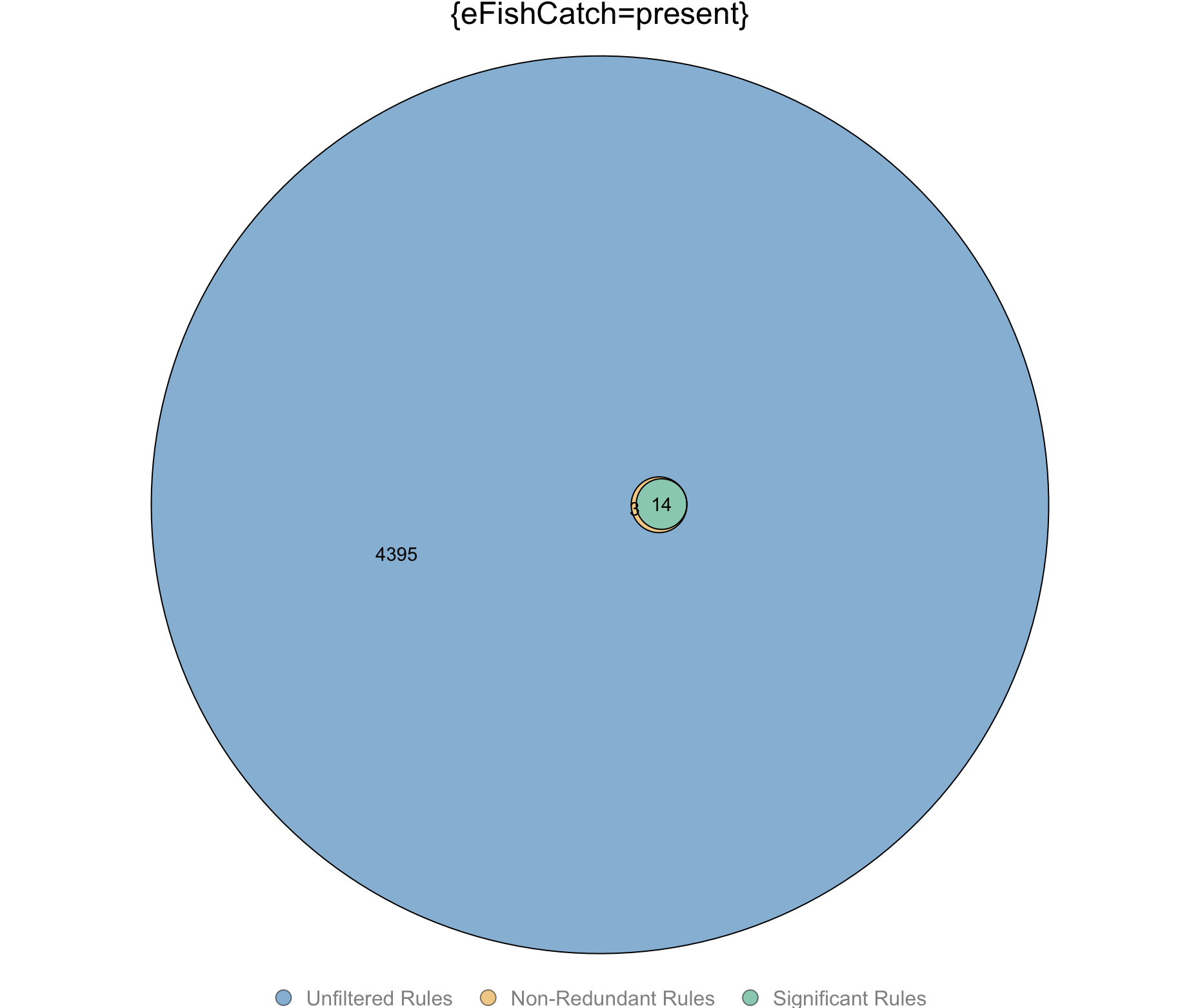

### SupplementaryFile7.png

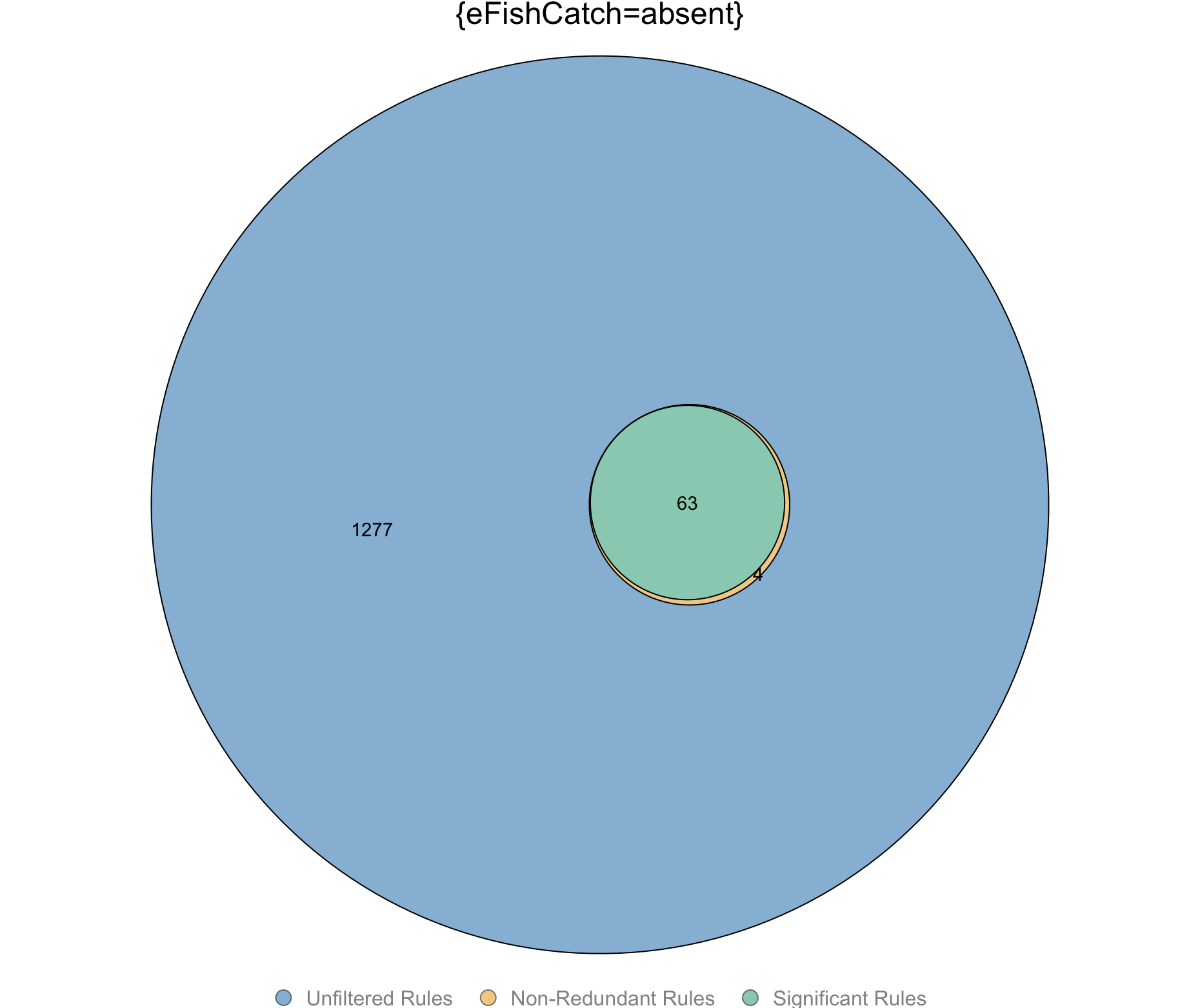

### SupplementaryFile8.png

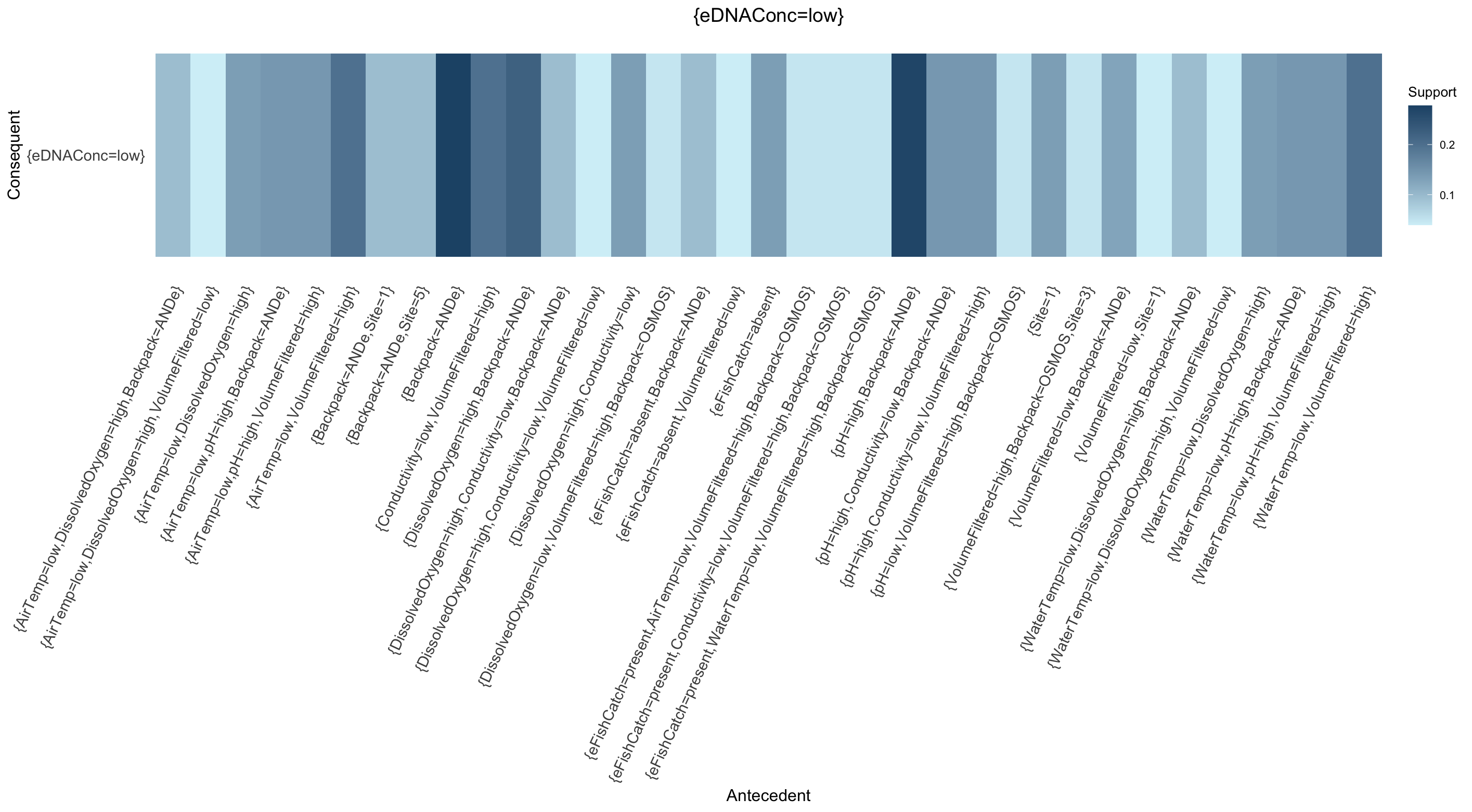

### SupplementaryFile9.png

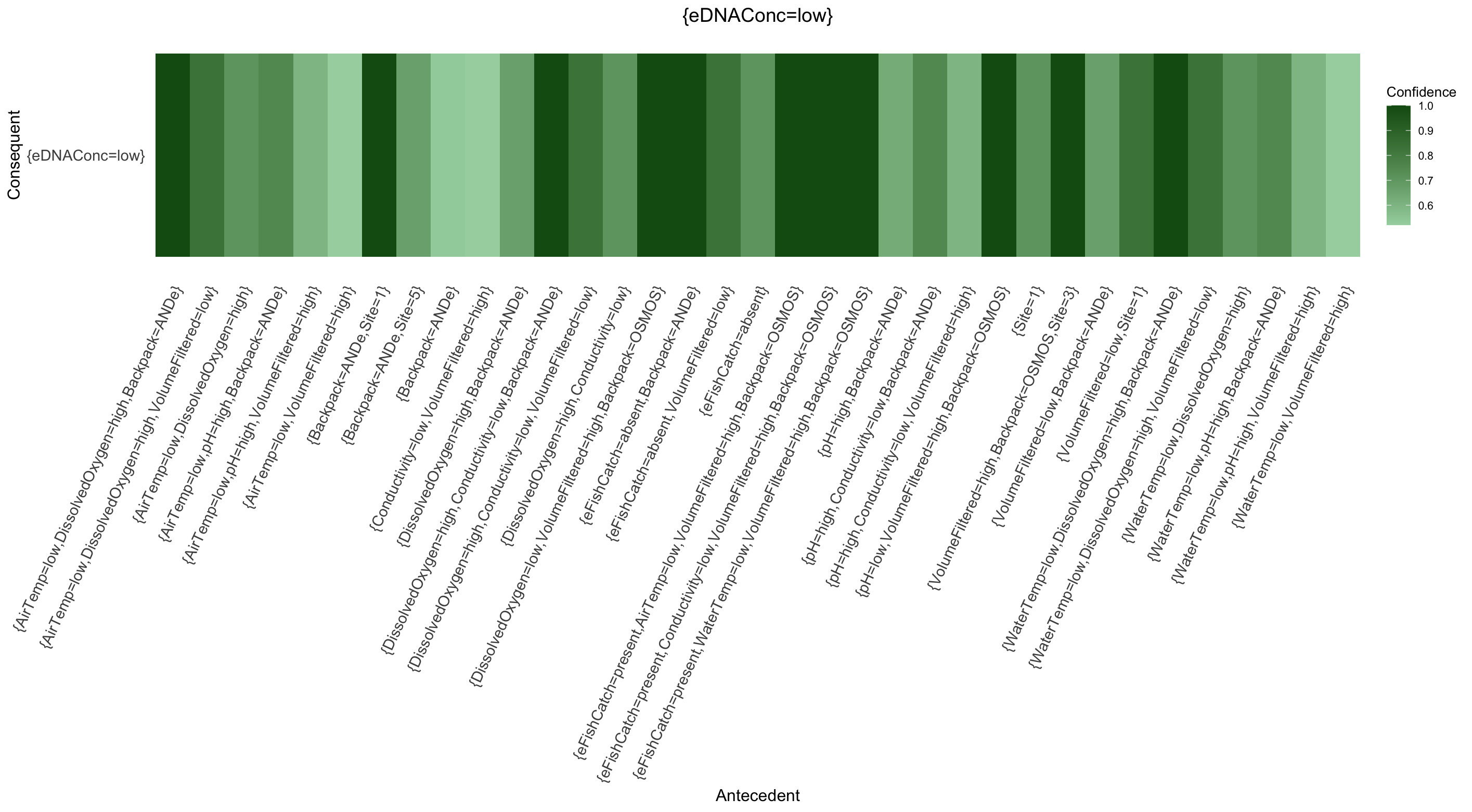

### SupplementaryFile10.png

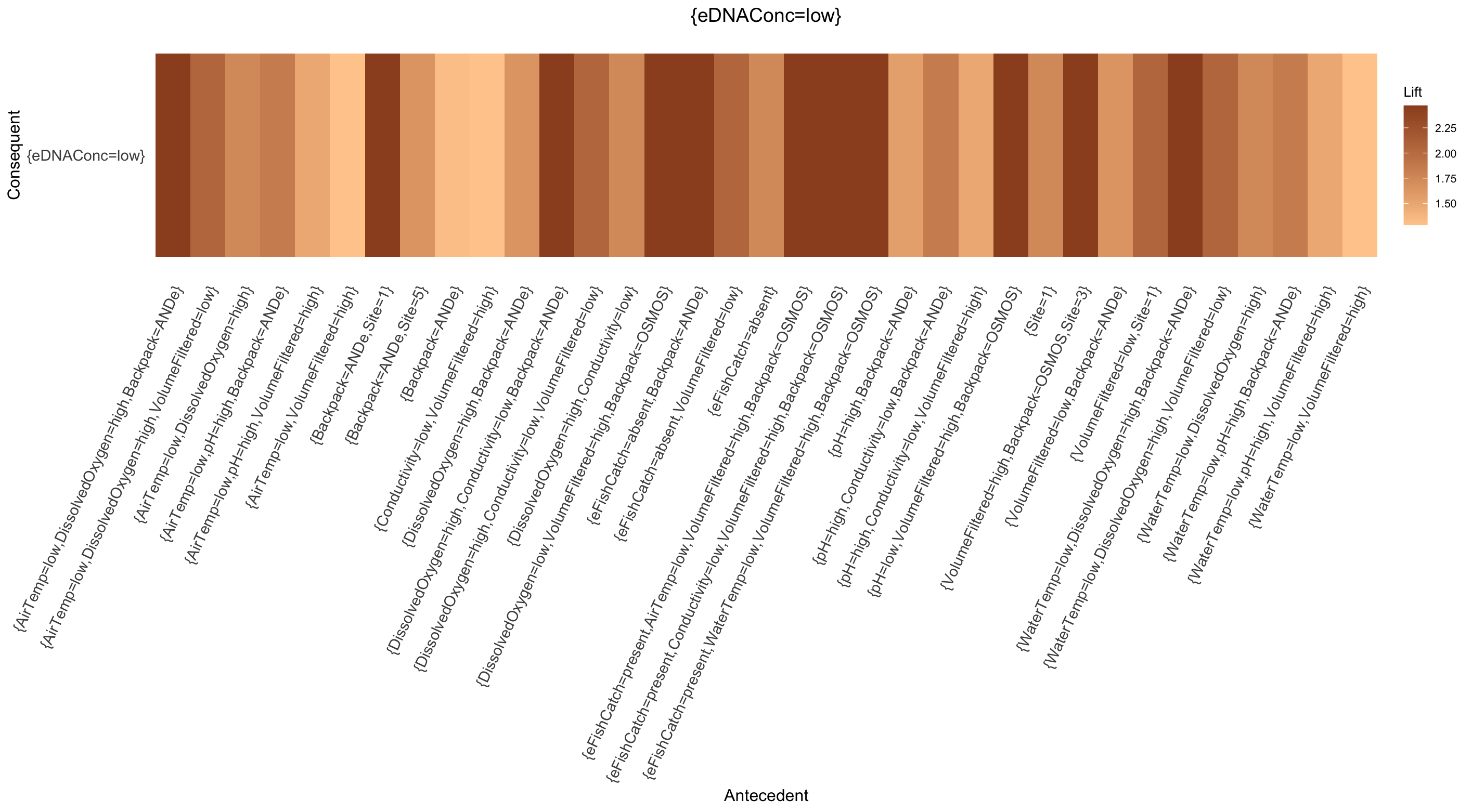

### SupplementaryFile11.png

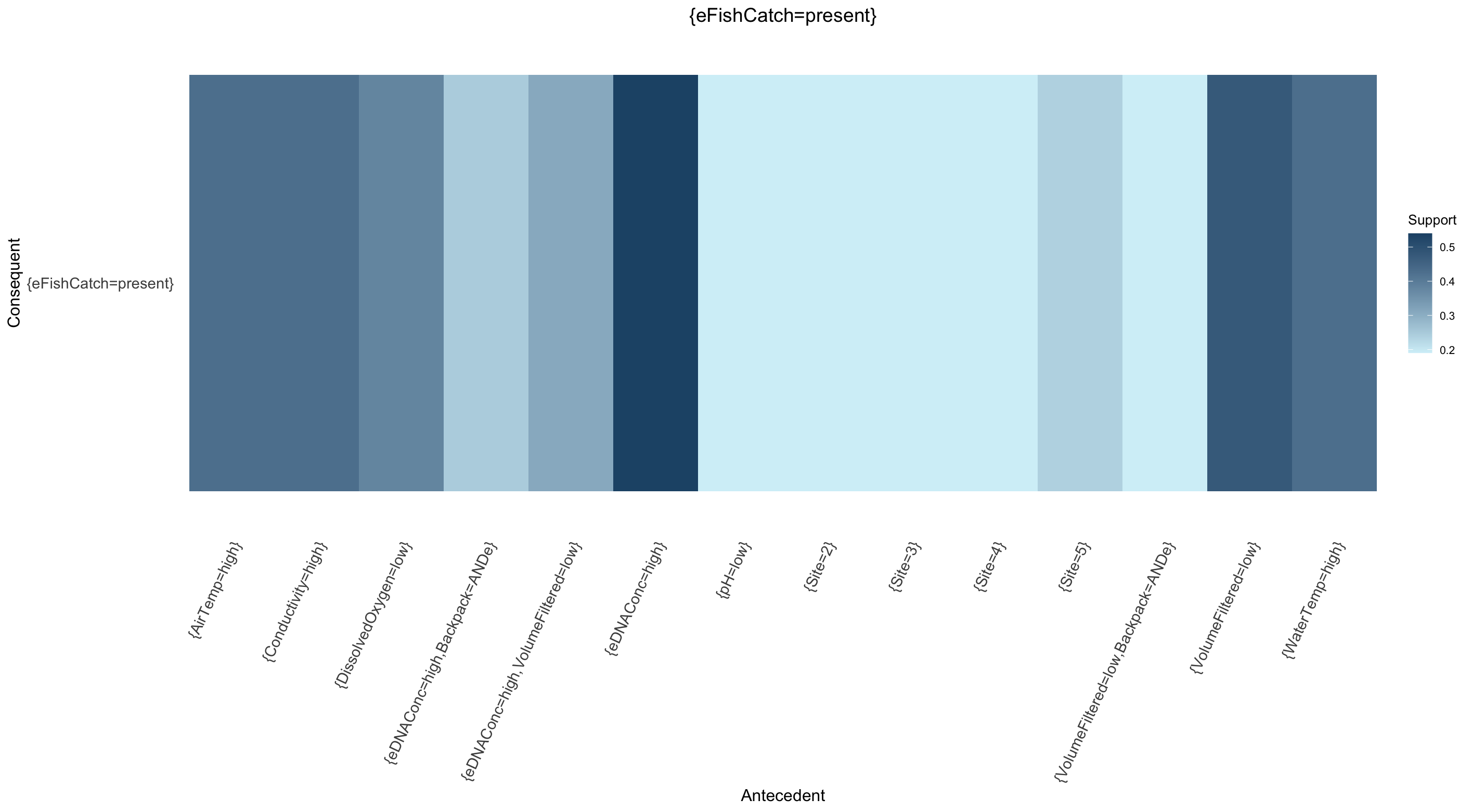

### SupplementaryFile12.png

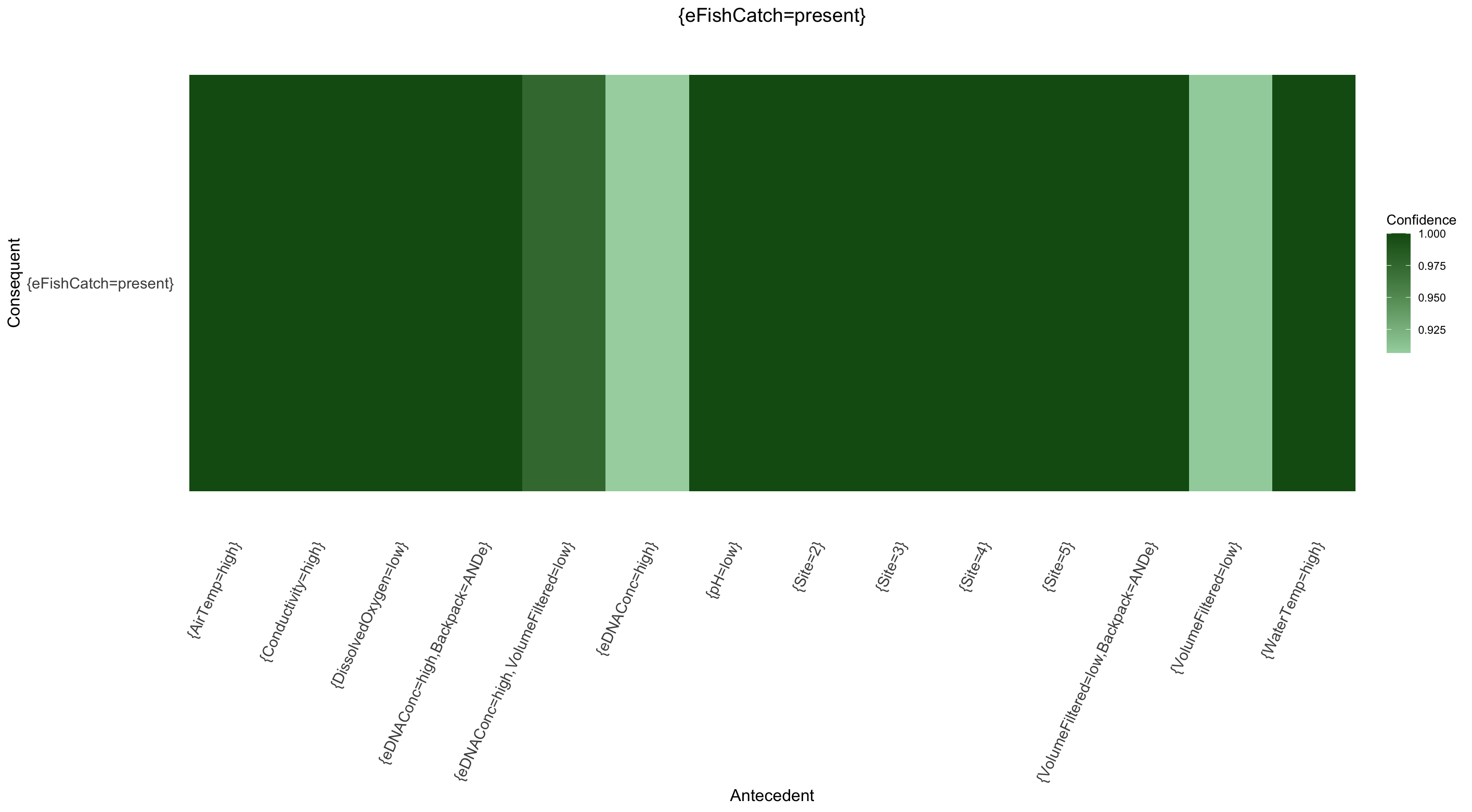

### SupplementaryFile13.png

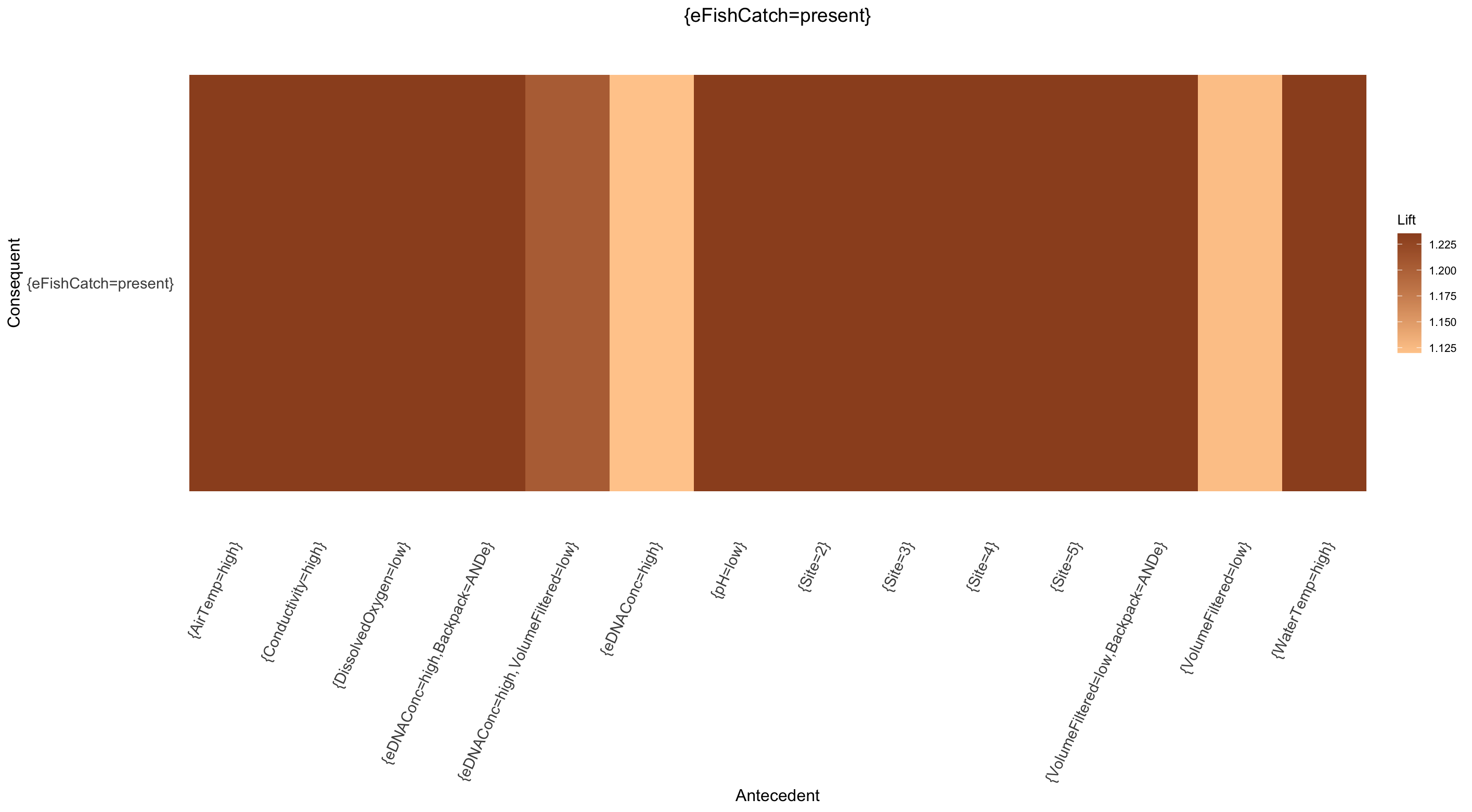

### SupplementaryFile14.png

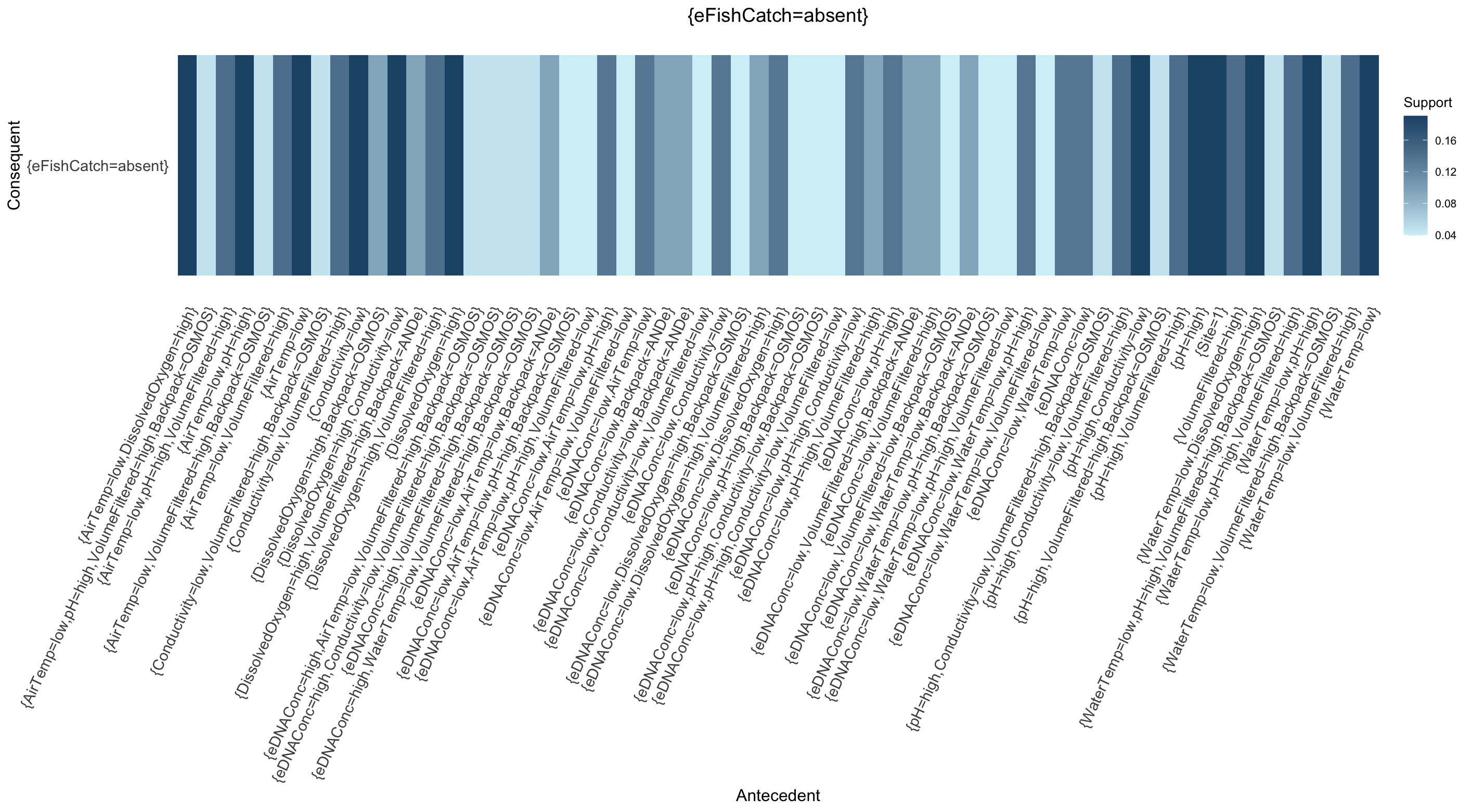

### SupplementaryFile15.png

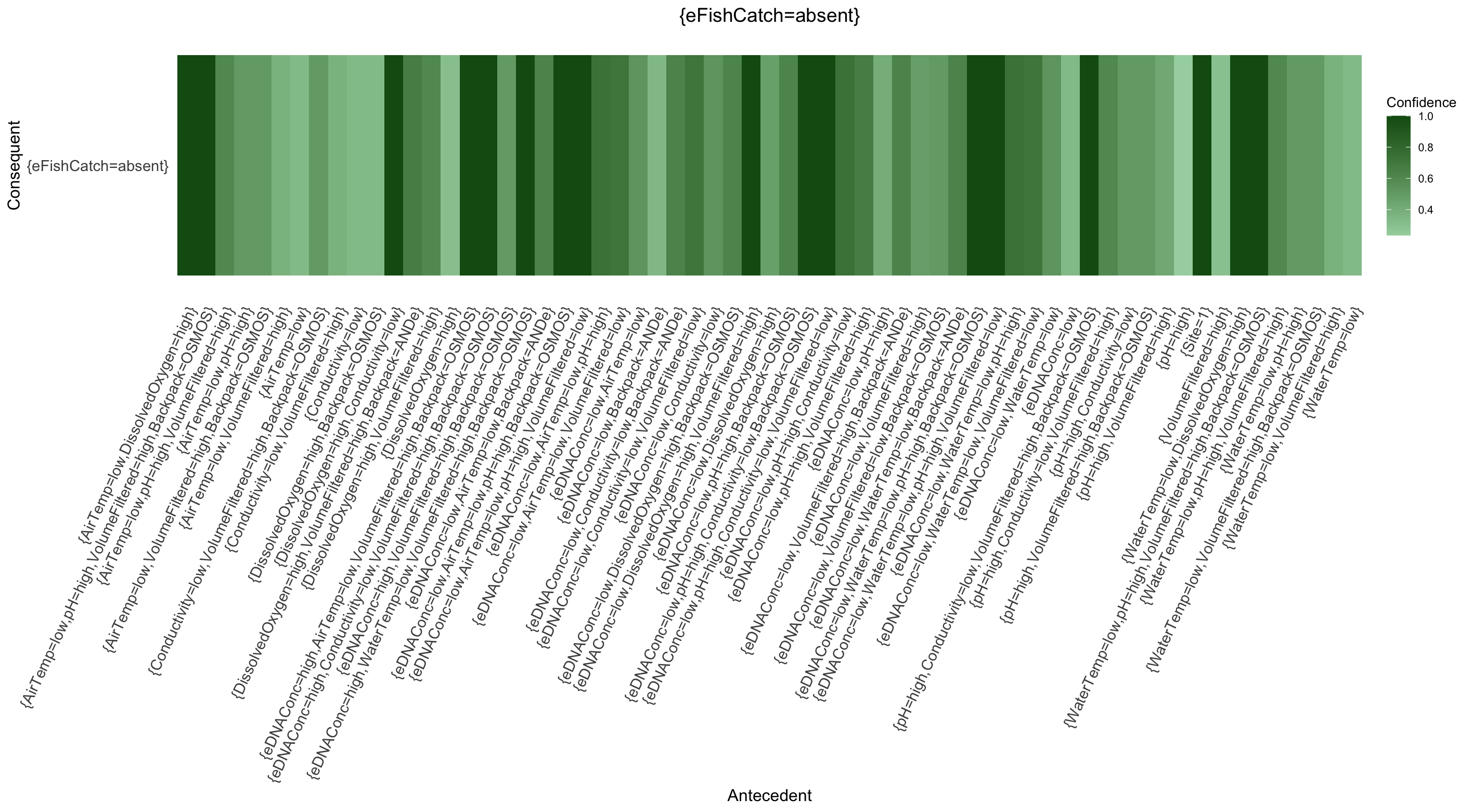

### SupplementaryFile16.png

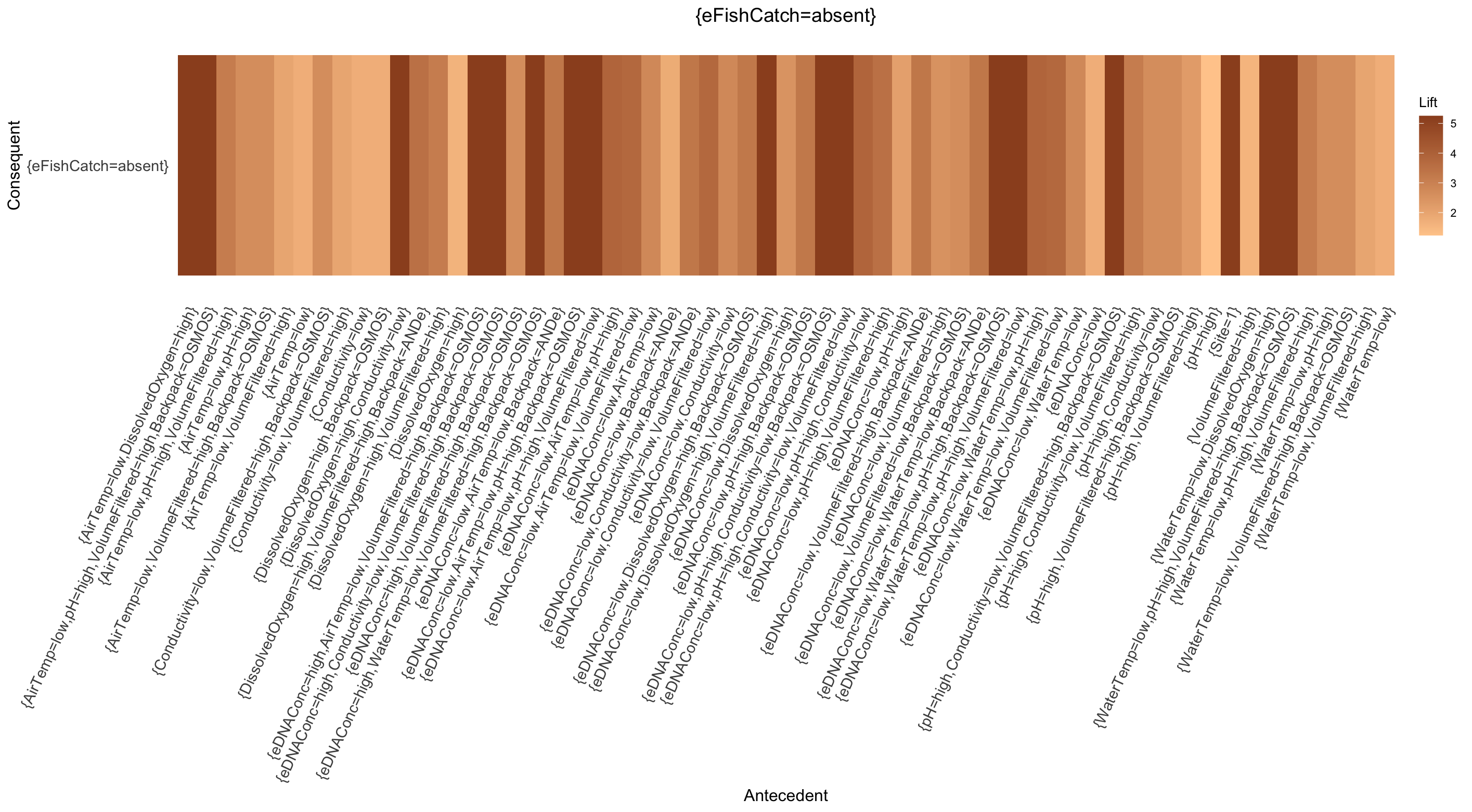
